# Combined KRAS^G12C^ and SOS1 inhibition enhances and extends the anti-tumor response in KRAS^G12C^-driven cancers by addressing intrinsic and acquired resistance

**DOI:** 10.1101/2023.01.23.525210

**Authors:** Venu Thatikonda, Hengyu Lu, Sabine Jurado, Kaja Kostyrko, Christopher A. Bristow, Karin Bosch, Ningping Feng, Sisi Gao, Daniel Gerlach, Michael Gmachl, Simone Lieb, Astrid Jeschko, Annette A. Machado, Ethan D. Marszalek, Mikhila Mahendra, Philipp A. Jaeger, Alexey Sorokin, Sandra Strauss, Francesca Trapani, Scott Kopetz, Christopher P. Vellano, Mark Petronczki, Norbert Kraut, Timothy P. Heffernan, Joseph R. Marszalek, Mark Pearson, Irene Waizenegger, Marco H. Hofmann

**Author notes:** These authors contributed equally (Venu Thatikonda, Hengyu Lu). **Corresponding Authors: Name**: Marco H. Hofmann, **Email**.

## Abstract

Efforts to improve the anti-tumor response to KRAS^G12C^ targeted therapy have benefited from leveraging combination approaches. Here, we compare the anti-tumor response induced by the SOS1-KRAS interaction inhibitor, BI-3406, combined with a KRAS^G12C^ inhibitor (KRAS^G12C^i) to those induced by KRAS^G12C^i alone or combined with SHP2 or EGFR inhibitors. In lung cancer and colorectal cancer (CRC) models, BI-3406 plus KRAS^G12C^i induces an anti-tumor response stronger than that observed with KRAS^G12C^i alone and comparable to those by the other combinations. This enhanced anti-tumor response is associated with a stronger and extended suppression of RAS-MAPK signaling. Importantly, BI-3406 plus KRAS^G12C^i treatment delays the emergence of acquired adagrasib resistance in both CRC and lung cancer models and is associated with re-establishment of anti-proliferative activity in KRAS^G12C^i-resistant CRC models. Our findings position KRAS^G12C^ plus SOS1 inhibition therapy as a promising strategy for treating both KRAS^G12C^-mutated tumors as well as for addressing acquired resistance to KRAS^G12C^i.

## Intro

Alterations in the KRAS gene are commonly found across all human cancers^1–3^, with the G12C mutant being one of the predominant KRAS variants occurring in non-small cell lung cancer (NSCLC; 13%) and colorectal cancer (CRC; 3%)^4,5^. Recent drug-discovery efforts^6,7^ have led to the clinical testing of multiple KRAS^G12C^-specific inhibitors (KRAS^G12C^i)^5,8^, with sotorasib (AMG 510)^9^ and adagrasib (MRTX849)^10^ demonstrating significant benefit in KRAS^G12C^-mutated NSCLC and CRC patients and both receiving approval for patients with KRAS^G12C^-mutated NSCLC^11,12^. While the success of these trials is impressive, disease progression occurs in the majority of cancer patients treated with KRAS^G12C^-inhibitors^9,11^. The rapid development of resistance to sotorasib or adagrasib can be recapitulated in preclinical studies^13–15^ therefore, intensive efforts are ongoing to identify approaches to improve the moderate rate and duration of response to KRAS^G12C^i monotherapy^5,13^.

KRAS functions as a molecular switch by cycling between inactive (GDP-bound) and active (GTP-bound) states, with exchange from the GDP-bound to the GTP-bound state regulated by guanine nucleotide exchange factors (GEFs) and intrinsic GTPase activity of RAS proteins enhanced by GTPase-activating proteins (GAPs)^16^. Active KRAS is crucial for controlling cell survival and proliferation through MAPK and PI3K pathway signaling, thus activating mutations in KRAS, which can result in the persistence of the GTP-bound state, enhance downstream signaling as well as promote rapid and uncontrolled cell growth^17^. The majority of current KRAS^G12C^i bind exclusively to GDP-bound KRAS^G12C^, locking it in its inactive state^18^. Preclinical and clinical studies have suggested that the activity of these “KRAS-off” inhibitors can be attenuated through mechanisms that reestablish RAS-MAPK signaling, either through shifting the cellular balance towards the increased activation of mutant or WT RAS, the acquisition of secondary KRAS mutations, and/or the acquisition of aberrations that enable upstream/downstream bypass events^5,11,13,14,19^.

Combinational approaches that co-target regulators of RAS GTP loading, such as receptor tyrosine kinases (RTKs) and downstream signaling intermediates, such as the phosphatase, SHP2, have been demonstrated to improve the efficacy and durability of KRAS^G12C^i response in pre-clinical NSCLC and CRC models^19–22^. Mechanistically, this approach works by addressing key feedback regulatory nodes and, at least in part, by shifting the balance of both mutant and WT KRAS to the GDP-bound state, thus enhancing the ability of KRAS^G12C^i to downregulate RAS-MAPK pathway signaling^19,21^. Based on these observations, both KRAS^G12C^i plus SHP2 inhibitors (SHP2i; e.g. TNO155) and KRAS^G12C^i plus EGFR antagonists (EGFRi; e.g. cetuximab or pantitumumab^23^) combination approaches have advanced to Phase I/II clinical trials in patients with KRAS^G12C^-mutated solid tumors^19,21,24,25^, with the latter being reported to achieve a response rate nearly double than that of KRAS^G12C^i monotherapy in patients with heavily pretreated KRAS^G12C^-mutated CRC^25,26^.

In pursuit of improving the efficacy and durability of KRAS^G12C^i, we have developed two specific and potent SOS1 inhibitors (SOS1i), BI-3406 and the clinical candidate BI 1701963, that target the SOS1-KRAS protein-protein interaction^27^. As a signaling intermediate following RTK activation, SOS1 directly controls the activation of the KRAS protein through its nucleotide exchange activity^28,29^. SOS1 acts as a key feedback node, where post-translational modification has been demonstrated to modulate the strength of signaling downstream of KRAS^27,28^. Inhibition of SOS1 thus enables enhanced and persistent maintenance of inactive GDP-bound KRAS^G12C^, similar to the effects observed under SHP2i^24^. Accordingly, the well-tolerated and prolonged anti-tumor response in KRAS-mutated solid tumor models treated with BI-3406 has been linked to the drug’s ability to obstruct the formation of GTP-loaded KRAS^27^.

Here, we hypothesize that SOS1 inhibition, by maintaining KRAS in the GDP-bound off state, may act as an effective combination partner with a covalent GDP-KRAS binding KRAS^G12C^i, such as adagrasib or sotorasib. As feedback re-activation of WT RAS may occur through multiple RTKs to drive resistance to KRAS^G12C^i, co-targeting a common node, such as SOS1, may induce a more durable response than co-targeting a single RTK, such as EGFR^21^. Further, by enhancing the activity of a KRAS^G12C^i, we anticipate that combination therapy may not only result in more pronounced effects in KRAS^G12C^i-naïve KRAS^G12C^-mutated tumors but also enable the re-establishment of tumor growth control in relapsed KRAS^G12C^-mutated tumors. To test our hypothesis, cellular growth inhibition, tumor response, and response duration elicited by KRAS^G12C^i plus BI-3406 treatment were assessed and compared to those elicited by KRAS^G12C^i alone or combined with TNO155 or cetuximab in KRAS^G12C^-driven NSCLC and CRC models. Our findings provide strong evidence that our KRAS^G12C^i plus SOS1i combination is a promising approach for treating KRAS^G12C^-mutated KRAS^G12C^i-naïve and resistant NSCLC and CRC.

## Results

### Adagrasib combined with BI-3406, TNO155, or cetuximab enhanced anti-proliferative responses in vitro and in vivo

We first conducted a high-throughput compound screen to broadly compare the anti-proliferative response of combining sotorasib (AMG 510) or adagrasib (MRTX849) with a panel (n=179) of small molecule inhibitors (**Suppl. Table 1**) in the KRAS^G12C^-driven NSCLC cell line, NCI-H2122. Potent synergistic anti-proliferative effects were observed with inhibitors blocking upstream activators of KRAS, including SOS1i (BI-3406), SHP2i (TNO155 and SHP099), and ErbB family inhibitors (lapatinib and afatinib) (**Fig. 1a**) as well as inhibitors that act downstream of KRAS, such as on the MAPK pathway (e.g. MEK inhibitor trametinib) or other pathways (e.g. PI3K inhibitor BYL719) (**Fig 1a**). Further testing of a representative compound subset against the KRAS^G12C^-driven CRC cell line, SW837, produced similar findings, with clear anti-proliferative responses but relatively lower synergy scores, that were likely due to the lower overall proliferation rates in these cells (**Fig. 1b**). Overall, our high-throughput screens found that treatment with an KRAS^G12C^i in combination with BI-3406, TNO155/SHP099, or lapatinib/afatinib consistently produced a synergistic anti-proliferative response.

**Figure 1.**
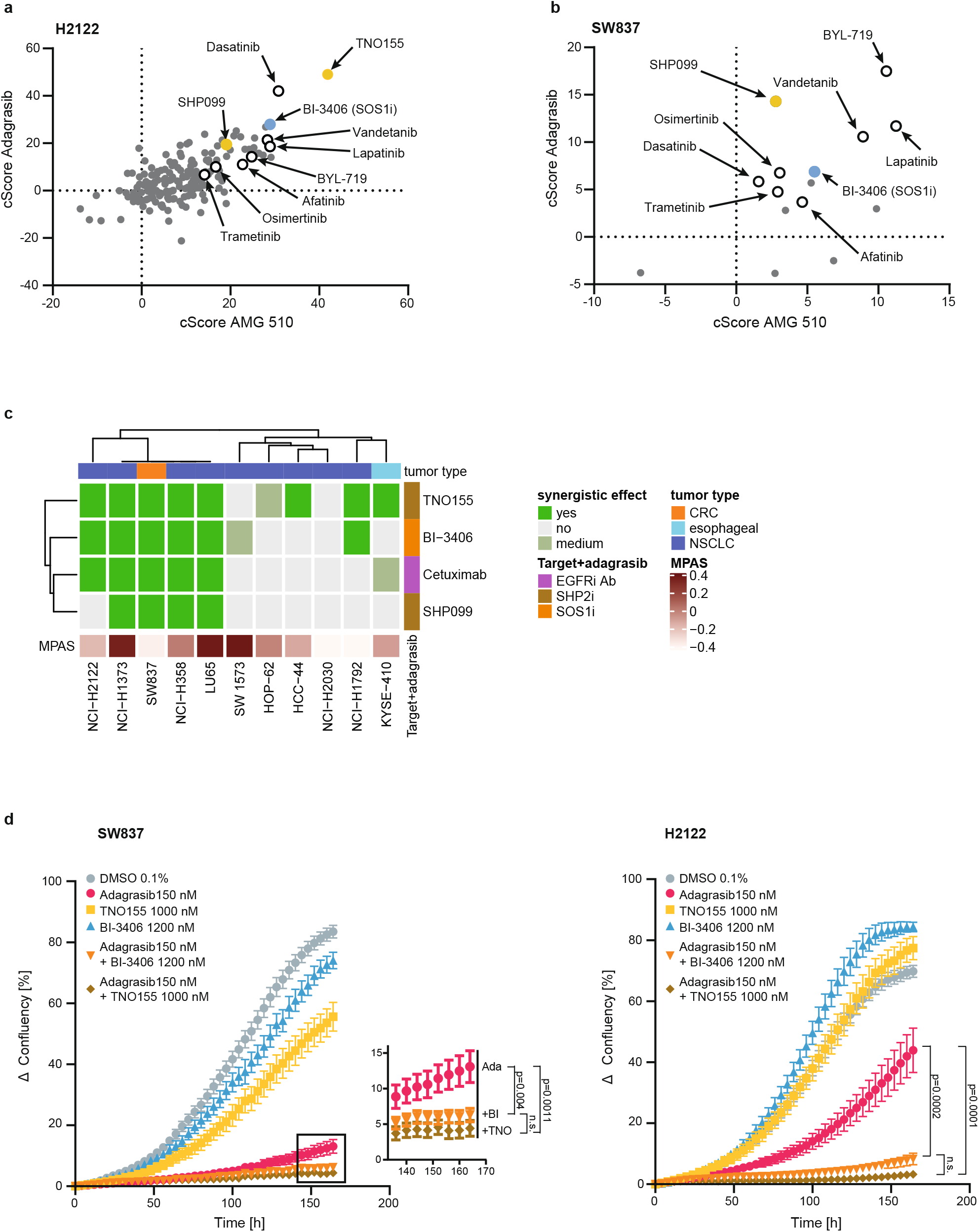
Efficacy of combination treatments in non-small cell lung cancer (NSCLC) and colorectal cancer (CRC) cell lines. **a**, Comparison of combination scores (cScore) indicating the response of the KRAS^G12C^ mutant NSCLC cell line, NCI-H2122, to the combination treatment of AMG 510 or adagrasib with 179 other small molecules for 72 hours. Positive scores indicate synergistic effects with scores >5-10 generally considered “clear synergy.” Empty or yellow/blue coloured circles highlight selected compounds for reference. **b**, Comparison of combination scores (cScore) indicating the response of KRAS^G12C^ mutant CRC cell line, SW837, to the combination treatment of AMG 510 or adagrasib with 15 other small molecules for 96 hours. Positive scores indicate synergistic effects with scores >5-10 generally considered “clear synergy”. Empty or yellow/blue coloured circles highlight selected compounds for reference. **c**, Response of indicated cell line to the combination treatment of adagrasib with an SOS1 inhibitor (BI-3406), a SHP2 inhibitor (TNO155 or SHP099), or an EGFR inhibitor (cetuximab). Cell lines are indicated as colorectal cancer (CRC; orange), esophageal (light blue), or non-small cell lung cancer (NSCLC; dark blue). MAPK Pathway Activity Score (MPAS) are indicated. **d**, Growth kinetic of NCI-H2122 (right) or SW837 (left) cells treated with indicated single or combination treatments, or vehicle control (N=3 for each curve). Y-axis indicates change in confluency relative to t=0 hours. Inset shows boxed area for clarity. Mean ± SEM shown. Data analyzed by one-way analysis of variance (ANOVA) with Tukey’s multiple comparisons test.

Findings from our high-throughput compound screens were strengthened using a panel of 11 KRAS^G12C^-driven tumor cell lines that were treated with adagrasib plus BI-3406, TNO155/SHP099, or cetuximab. The strongest synergistic anti-proliferative effects were observed upon treatment with adagrasib plus BI-3406 (6/11 lines) or TNO155 (8/11 lines) (**Fig 1c**). Notably, the synergistic effect on proliferation upon combination of adagrasib plus BI-3406 was observed in the same cell lines where synergy was seen upon combination treatment with TNO155 (**Fig 1c)**. Additionally, strong synergistic anti-proliferative effects were observed upon treatment with adagrasib plus cetuximab (5/11 lines) or another SHP2i, SHP099 (4/11 lines) (**Fig. 1c**). To further confirm these findings, we analyzed the effect of adagrasib plus BI-3406 or TNO155 on the *in vitro* growth kinetics of SW837 and NCI-H2122 cell lines over seven days (**Fig 1d**). In both cell lines, treatment with adagrasib alone resulted in a strong initial anti-proliferative effect that attenuated over time. In contrast, adagrasib plus BI-3406 or TNO155 resulted in a more profound and durable anti-proliferative effect on cell confluency (**Fig. 1d)**.

The enhanced anti-tumor benefit gained from adagrasib plus BI-3406, TNO155, or cetuximab therapy was further confirmed *in vivo* using both the NCI-H2122 and SW837 cell line-derived (CDX) as well as two CRC patient-derived xenograft (PDX) models. Adagrasib alone resulted in modest tumor growth inhibition (TGI of 83% on Day 16) in the NCI-H2122 model, with an initial phase of tumor stasis followed by outgrowth (**Fig 2a**), suggesting onset of resistance. When compared to adagrasib monotherapy, treatment with adagrasib plus BI-3406 (TGI of 106%, average tumor volume (TV) change from baseline of −18% on Day 16) or TNO155 (TGI of 104%, average TV change from baseline −13% on Day 16) resulted in significantly enhanced growth inhibition in the NCI-H2122 model (p=0.0222, 5/6 tumors in regression from BI-3406 co-administration; p=0.0189, 5/7 tumors in regression from TNO155 combination) (**Fig 2a**). Tumor outgrowth, albeit slow, was still observed in the NCI-H2122 model upon combination treatment over the duration of the experiment (**Fig 2a**). In the SW837 xenograft model, the combination of adagrasib plus BI-3406 (TGI of 119%, average TV change from baseline of −67.8% on Day 42) or cetuximab (TGI of 121% on Day 42, average TV change from baseline of −74.4%) resulted in enhanced efficacy compared to adagrasib alone (TGI of 108%, average TV change from baseline of −35%), with combinations inducing deeper regressions in all tumors (**Fig 2b**). Similarly, in CRC PDX models F3008 and B8032, treatment with adagrasib alone induced modest anti-tumor activity, with tumor outgrowth observed around Day 20 post-treatment, while adagrasib plus either BI-3406 or cetuximab resulted in tumor regression throughout the experimental observation period (**Fig 2c,d**). Overall, the *in vivo* anti-tumor activity induced by adagrasib plus BI-3406 treatment was comparable to that induced by adagrasib plus TNO155 or cetuximab in NSCLC and CRC models.

**Figure 2.**
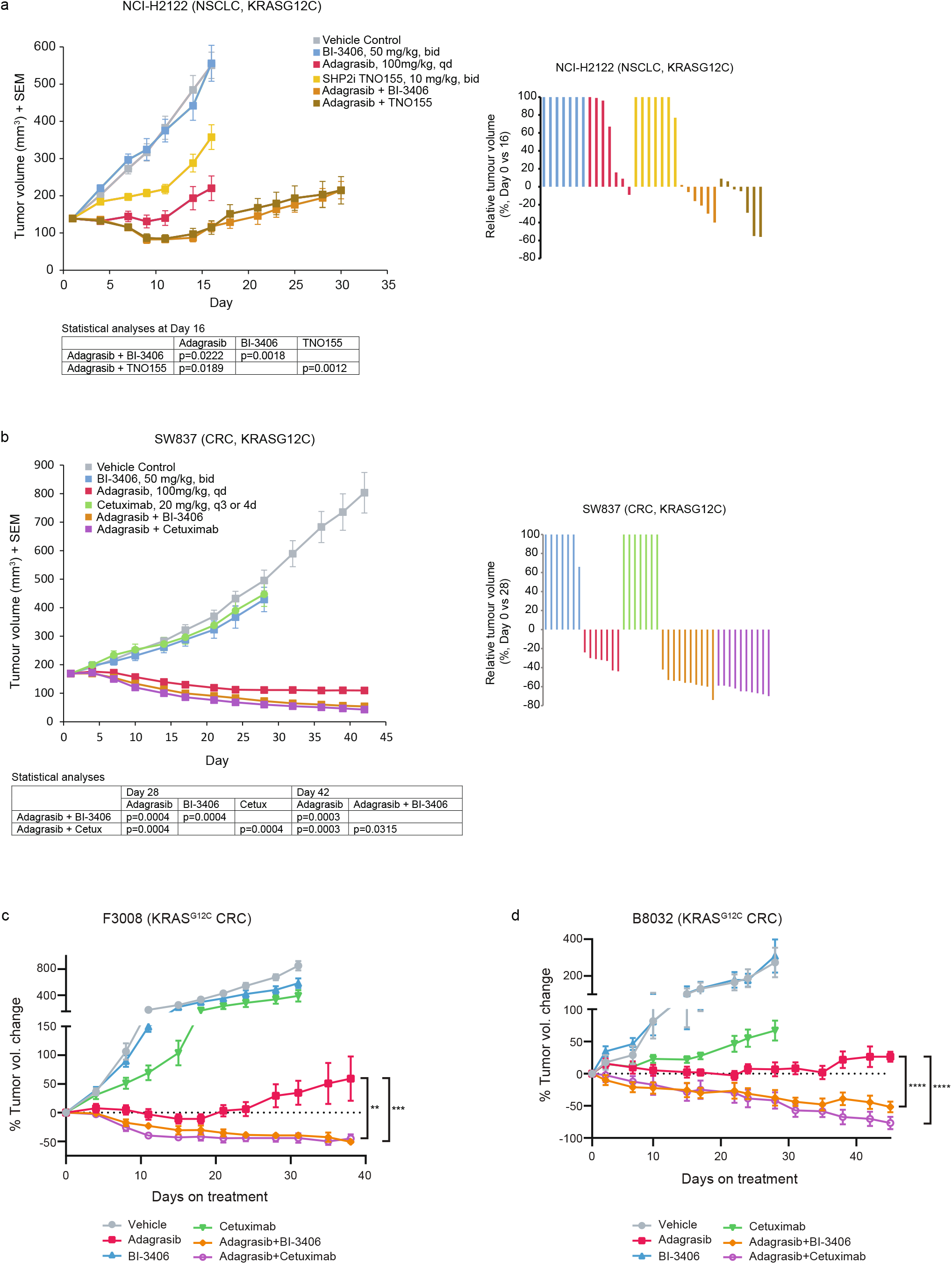
Efficacy of combination treatments in non-small cell lung cancer (NSCLC) and colorectal cancer (CRC) xenograft models. **a**, (*left*) Tumor volumes of mice injected subcutaneously with NCI-H2122 cells. Mice were treated 5 days of each week orally either with control vehicle, BI-3406 (50mg/kg; bid, with a delta of 6 hours), TNO155 (10mg/kg, bid), adagrasib (100mg/kg, qd) for 16 days, or with the combination of adagrasib plus BI-3406 or TNO155 for 30 days. n=7 animals/group. Data analyzed by one-sided Mann-Whitney-Wilcoxon U-tests adjusted for multiple comparisons according to the Bonferroni-Holm Method within each subtopic. Mean ± SEM shown. (*right*) Relative NCI-H2122 tumor volumes are indicated as percent change from baseline at Day 16. Values smaller than zero percent indicate tumor regressions. **b**, (*left*) Tumor volumes of mice injected subcutaneously with SW837 cells. Mice were treated 5 days of each week orally either with control vehicle, BI-3406 (50mg/kg; bid; with a delta of 6 hours), (100mg/kg, qd), cetuximab (20mg/kg, q3 or 4d, intraperitoneal), or adagrasib (100mg/kg for 28 days, qd), or with the combination of adagrasib plus cetuximab or BI-3406 for 42 days. n=7/group for monotherapies; n=10/group for combination therapies. Data analyzed by one-sided Mann-Whitney-Wilcoxon U-tests adjusted for multiple comparisons according to the Bonferroni-Holm Method within each subtopic. Mean ± SEM shown. (*right*) Relative SW837 tumor volumes are indicated as percent change from baseline at Day 28. Values smaller than zero percent indicate tumor regressions. **c**, Change in tumor volume in NSG mice implanted with KRASG12C-driven F3008 patient-derived xenograft (PDX) fragments and treated with vehicle, adagrasib (100 mg/kg, daily), BI-3406 (50mg/kg, twice daily), cetuximab (15 mg/kg, twice weekly), or the combination of adagrasib plus BI-3406 or cetuximab. n=8/group. Data analyzed by paired Student’s T-test; **p < 0.01; ***p < 0.001. Mean ± SEM shown. **d**, Change in tumor volume in nude mice implanted with KRASG12C-driven B8032 PDX fragments and treated with vehicle, adagrasib (100 mg/kg, daily), BI-3406 (50mg/kg, twice daily), cetuximab (20 mg/kg, twice weekly), or the combination of adagrasib plus BI-3406 or cetuximab. N = 5/group. Data analyzed by paired Student’s T-test; ***p < 0.001. Mean ± SEM shown.

Together, our findings demonstrate that anti-tumor effects achieved by adagrasib plus BI-3406 therapy are similar to those induced by adagrasib plus TN0155 or cetuximab, both currently under clinical evaluation. These preclinical findings have contributed to launching the clinical assessment of KRAS^G12C^i plus SOS1i (BI 1701963) combination therapy (NCT04185883 and NCT04973163).

### Combination therapy induces greater inhibition of the RAS-MAPK pathway and cell proliferation than monotherapy in vitro

Oncogenic activity is enabled through interactions with the GTP-bound (active) form of mutated KRAS, with mediators, such as RAF or PI3K, regulating activation of downstream signaling intermediates, such as the RAS-MAPK pathway^5^. Modulation of KRAS-GTP levels by adagrasib plus BI-3406 treatment was thus assessed and compared to that induced by monotherapy or other combinatory approaches. Using a G-LISA assay, adagrasib plus BI-3406 treatment was demonstrated to further reduce RAS-GTP levels, particularly at 7- and 24-hours following treatment, when compared to the effects induced by adagrasib or BI-3406 monotherapy in the NCI-H2122 cell line (**Fig 3a**). These findings agree with the expected activity of a SOS1 KRAS-GEF inhibitor that prevents the exchange of GTP for GDP and, in the presence of adagrasib, traps KRAS in the GDP bound inactive form.

**Figure 3.**
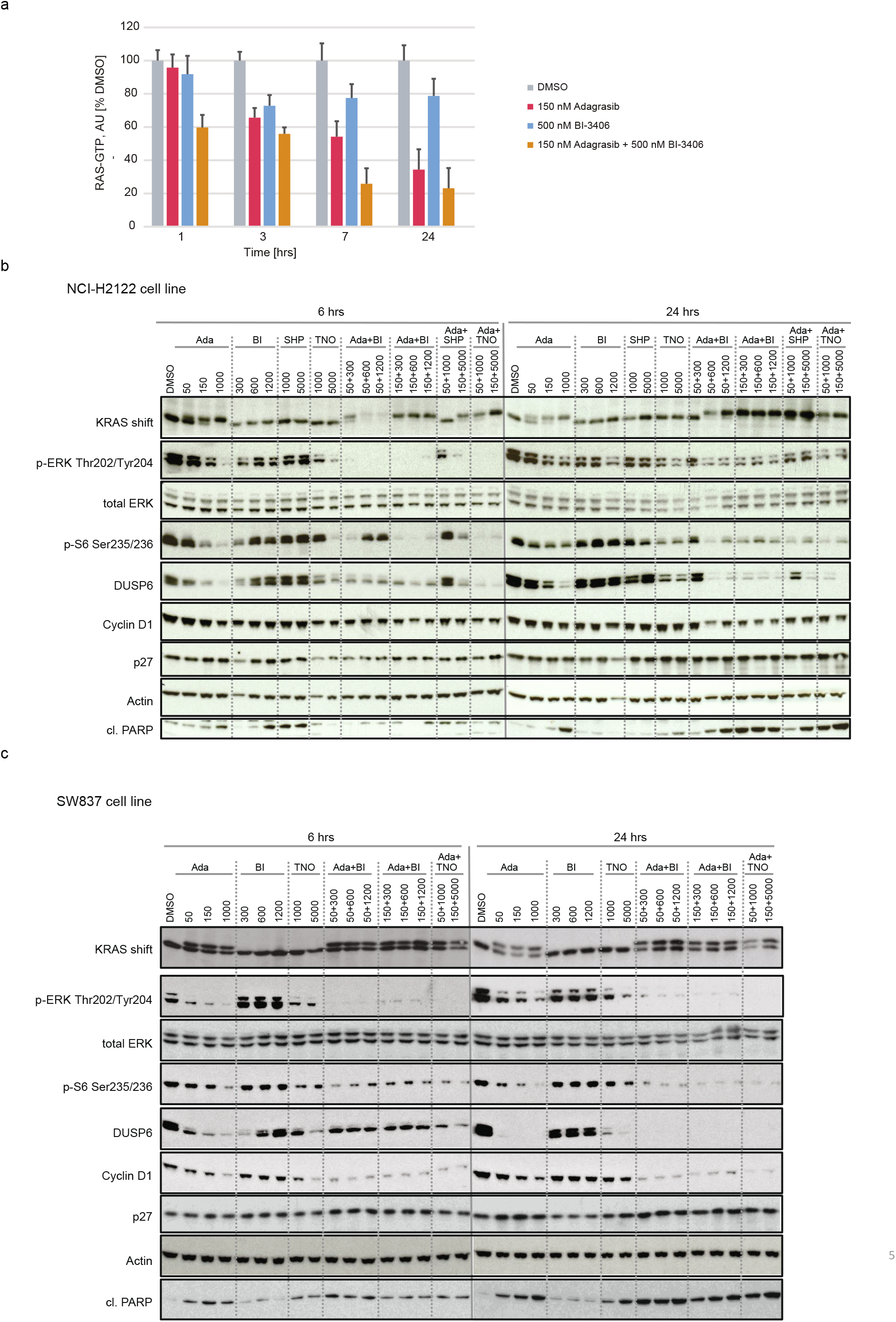
Modulation of RAS-MAPK signaling by adagrasib monotherapy or combination treatments in non-small cell lung cancer (NSCLC) and colorectal cancer (CRC) cell lines. **a**, GLISA plate analysis of RAS GTP levels across indicated time points in KRAS^G12C^ mutant lung cancer (NCI-H2122) cells treated with DMSO, adagrasib (150 nM), BI-3406 (500 nM), or the combination of adagrasib plus BI-3406. Data shown as % of the DMSO control at each indicated time point; error bars = SD. n=2. **b**, Western blot analysis of NCI-H2122 cells treated with adagrasib, BI-3406, TNO155, or SHP099 alone, or adagrasib combined with BI-3406, SHP099, or TNO155 at indicated concentrations for either 6 or 24 hours. KRAS shift indicates covalent binding of compound to KRAS^G12C^ resulting in a slower migrating form of KRAS. β-Actin served as the loading control. **c**, Western blot analysis of the KRAS^G12C-^mutant colon cancer cell line, SW837, treated with adagrasib, BI-3406, TNO155, or SHP099 alone, or adagrasib combined with BI-3406, SHP099, or TNO155 at indicated concentrations for either 6 or 24 hours. KRAS shift indicates covalent binding of compound to KRAS^G12C^, resulting in a slower migrating form of KRAS. β-Actin served as the loading control. Of note, to detect total ERK and cl. PARP, only 15 μg total protein per lane was loaded. **b-c**, Ada: adagrasib; BI: BI-3406; SHP: SHP099; TNO: TNO155

The impact of combination therapies on KRAS downstream signaling was then investigated. Pharmacodynamic markers of the RAS-MAPK pathway and changes in cellular phenotype after 6 and 24 hours of treatment were evaluated following single-agent or combination treatment with adagrasib, BI-3406, and/or TNO155/SHP099. Study concentrations were matched to clinically relevant plasma exposures that were determined through clinical data^9–11,30^, or through preclinical and investigational new drug studies that leveraged several cancer models. After 6 hours of treatment, adagrasib plus BI-3406 or TNO155/SHP099 induced stronger inhibitory effects on MAPK and PI3K signaling, as evidenced by reductions in pERK and p-S6, compared to those induced by adagrasib, TNO155, or BI-3406 alone in both cell lines (**Fig 3b, c**). Further, after 24 hours of treatment, combination approaches or high doses of adagrasib monotherapy achieved a sustained inhibition of downstream MAPK signaling effectors, such as the negative regulator, DUSP6, as well as increased cleaved PARP levels in both cell lines (**Fig 3b, c)**. Of note, a rebound of p-ERK and p-S6 levels was observed in cells treated with adagrasib monotherapy, combination therapy shown with either BI-3406 or SHP2i showed less of re-bound signals (**Fig. 3b, c**). These effects were associated with a significant reduction in Cyclin D1 levels in SW837 cells, with a more modest effect in NCI-H2122 cells, after 24 hours of combination treatment (**Fig 3b, c**). Together, these findings suggest that combination approaches were capable of inducing cell cycle arrest in NCI-H2122 and SW837 cells *in vitro*, with relatively stronger anti-proliferative effects on SW837 cells as seen with the increased PARP cleavage (**Fig 3b, c**).

Overall, our findings demonstrated that combining adagrasib with either BI-3406 or TNO155/SHP099 induced more profound modulation of MAPK pathway signaling and proliferative biomarkers than single-agent approaches. These findings correlated with the enhanced anti-tumor activity observed when these drugs were combined.

### Inhibition of the RAS-MAPK pathway is associated with the anti-proliferative effects of KRAS^G12C^i combination therapy in vivo

We next assessed treatment-induced inhibitory effects on pharmacodynamic biomarkers of the RAS-MAPK pathway *in vivo*. To this end, tumor-bearing mice were treated with either single-agent or combination treatments for 7 consecutive days, with samples collected at 4 and 48 hours after the last dose for further analysis.

Differential gene expression analysis was first conducted on NCI-H2122 xenograft models treated with adagrasib alone or combined with BI-3406 or TNO155, and SW837 xenograft models treated with adagrasib alone or combined with BI-3406 or cetuximab. In the NCI-H2122 xenograft model, both combination treatments induced a stronger overall transcriptomic change, when compared to that induced by adagrasib alone at 4 hours post-last dose. Specifically, 578 and 547 genes were modulated upon treatment with adagrasib plus BI-3406 or TNO155, respectively, as compared to the 217 genes deregulated upon adagrasib monotherapy (**Fig 4a**). Similarly, in SW837 xenograft models, 1,233 and 1,307 genes were differentially regulated upon treatment with adagrasib plus BI-3406 or cetuximab, respectively, and 629 genes were deregulated upon adagrasib monotherapy (**Extended Data Fig 1a**). Of note, the largest overlap of deregulated genes was identified between adagrasib plus BI-3406 and adagrasib plus cetuximab in SW837 xenograft tumors, suggesting that these combination strategies induced similar transcriptomic modulation (**Extended Data Fig 1a, Supp. Table 3)**.

**Figure 4.**
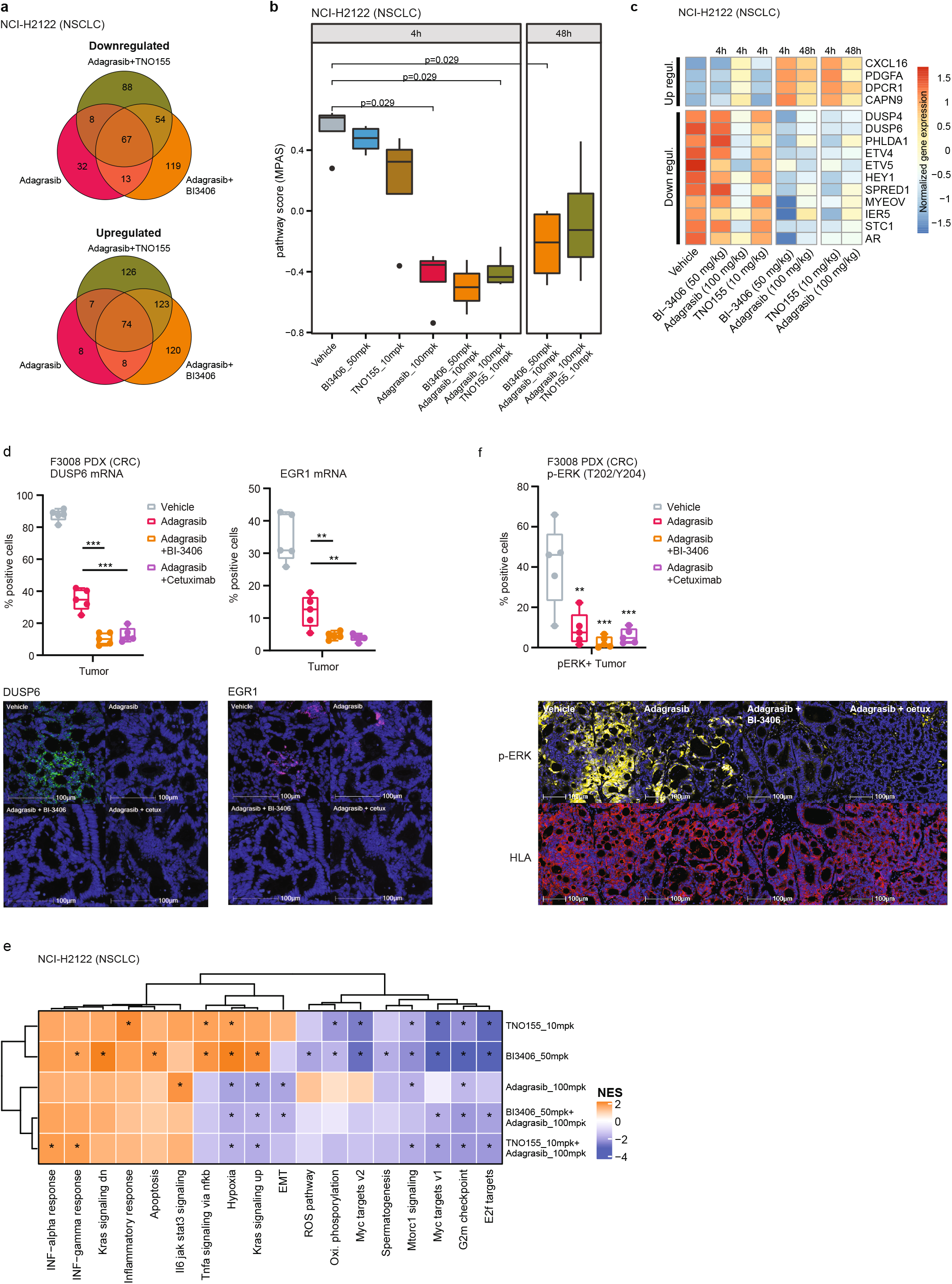
Differential gene expression induced by single-agent or combination treatments in vivo. All treatments were administered for 7 consecutive days, and samples were collected after the last dose. **a**, Venn diagram showing the overlap of upregulated (top) or downregulated (bottom) differentially expressed genes in the KRAS^G12C^-mutant non-small cell lung cancer xenograft model (NCI-H2122) xenograft models treated with adagrasib (100 mg/kg) alone or combined with BI-3406 (50 mg/kg) or TNO155 (10 mg/kg). **b**, Modulation of overall MAPK pathway activity, as indicated by the MAPK Pathway Activity Score (MPAS), in NCI-H2122 xenograft models treated with indicated monotherapies or combination treatments at 4 or 48 hours post-last doses. N=2-3. Boxplots show low and upper quartiles and median line is indicates. Whiskers: 1.5 x interquartile range. Data analyzed by two-sided Wilcoxon rank sum test. **c**, Differential modulation of select MAPK pathway genes in NCI-H2122 xenograft models treated with indicated single-agent or combination treatments. Samples were collected at 4 or 48 hours post last dose. **d**, (*top*) Percentage of positive tumor cells expressing *DUSP6* or *EGR1* mRNA analyzed by RNAscope in F3008 colorectal cancer (CRC) patient-derived xenograft (PDX) models treated with vehicle, adagrasib (100mpk, QD), adagrasib plus BI-3406 (50mpk, BID), or adagrasib plus cetuximab (15mpk, twice weekly) for 5 days. Tumors were collected 4 hours after the last dose. N=5/group. Boxplots show low and upper quartiles and median line is indicates. Whiskers: 1.5 x interquartile range. Data analyzed by two-sided Student’s t-test, **p<0.01, ***p<0.001. (*bottom*) Representative images from multiplex immunofluorescence analysis of DUSP6 and EGR1 in tumor tissue from models described in (*top*). Scale bar = 100 μm. **e**, Gene set enrichment analysis heatmaps show differentially modulated pathways in NCI-H2122 xenograft models treated with indicated single-agent or combination therapies. **f**, (*top*) Percentage of positive tumor cells expressing phospho-ERK as analyzed by multiplex immunofluorescence in the same tumor tissues as described in **d**. N=5/group. Boxplots show low and upper quartiles and median line is indicates. Whiskers: 1.5 × interquartile range. Comparisons to vehicle analyzed by two-sided Student’s t-test; **p<0.01, ***p<0.001. (*bottom*) Representative images from multiplex immunofluorescence analysis of p-ERK and HLA in tumor tissue from models described in (*top*). Scale bar = 100 μm.

We then quantified the overall MAPK pathway activity of individual tumors by conducting MPAS (MAPK Pathway Activity Score) analysis^31^ on transcriptomic data as well as in depth investigation of the modulation of key MAPK regulated genes. In NCI-H2122 xenograft tumors, treatment with adagrasib plus BI-3406 (median MPAS score = −0.502 at 4 hours post-last dose) or TNO155 (median MPAS score = −0.435 at 4 hours post-last dose) achieved more pronounced overall downregulation of MAPK activity than that observed in response to adagrasib alone (p=0.029 for all comparisons; median MPAS score = −0.35 at 4 hours post-last dose) (**Fig 4b**). Consistently, both combination treatments also achieved greater downregulation of genes regulated by the MAPK pathway (e.g., DUSP6) at 4 and 48 hours post-last dose (**Fig 4c, Supp Table 2**). In SW837 xenograft tumors, adagrasib alone (median MPAS score = −0.57) or combined with BI-3406 (p=0.029; median MPAS score = −0.49) or cetuximab (p=0.016; median MPAS score = −0.68) all achieved downregulation of overall MAPK activity at 4 hours post-last dose (**Extended Data Fig 1b, c**). However, only combination approaches were able to sustain downregulation of MAPK pathway activity, including suppressing expression of CCND1 (log 2 change compared to vehicle control: −1.8 fold) and DUSP6 (log 2 change compared to vehicle control: −2.4 fold), up to 48 hours post-last dose (**Extended Data Fig 1b, c; Supp. Table 3**). These findings were confirmed in the F3008 CRC PDX model using an RNAScope *in situ* hybridization assay, where treatment with adagrasib plus BI-3406 or cetuximab resulted in a greater downregulation of DUSP6 and EGR1 than that observed in response to monotherapy (**Fig 4d**). Of note, the strength of treatment-induced MAPK pathway downregulation (**Extended Data Fig 1b**) and gene modulation (**Extended Data Fig 1a**) were similar in tumors derived from SW837 models treated with adagrasib plus BI-3406 or cetuximab.

Downregulation of RAS-MAPK signaling likely mediated the enhanced anti-proliferative effects induced by combination approaches^5^ in NCI-H2122 and SW837 cell lines (**Fig 1, Fig 3b,c**), xenograft models (**Fig 2a,b**), and CRC PDX models (**Fig 2c,d**). To investigate this further, gene set enrichment analysis (GSEA) was used to identify the biological processes enriched after treating NCI-H2122 xenograft tumors with adagrasib or BI-3406 alone or adagrasib plus BI-3406 or TNO155, as well as SW837 xenograft tumors with adagrasib or cetuximab alone or adagrasib plusBI-3406 or cetuximab. In NCI-H2122 xenograft tumors, pathways associated with cell growth and survival, epithelial to mesenchymal transition (EMT), targets of oncogenic transcription factor *MYC*, as well as pathways up-regulated by active *KRAS* were observed to be consistently downregulated in response to combination therapies (*FDR < 0.01*) (**Fig 4e**). Similarly, in SW837 xenograft tumors, pathways associated with cell growth and survival, as well as *MYC* target pathways, were downregulated in response to single-agent and combination treatments (**Extended Data Fig 2**). Interestingly, immunohistochemistry (IHC) analysis of NCI-H2122 tumors revealed that combination with BI-3406, but not TNO155, induced a significantly stronger downregulation of both cell proliferation biomarker, Ki-67, and cell growth biomarker, phospho-ERK (p=0.0318 and p=0.014 for p-ERK and Ki67, respectively, at 4 hours post-last dose) when compared to that induced by adagrasib monotherapy (**Extended Data Fig 3a**). In comparison, IHC analyses of SW837 xenograft tumors demonstrated that, when compared to adagrasib monotherapy, both adagrasib plus BI-3406 (p=0.008 and 0.079 at 4 and 48 post-last dose, respectively) or cetuximab (p=0.008 at both 4 and 48 hours post-last dose) significantly downregulated Ki-67 to similar levels (**Extended Data Fig 3b)**. However, all therapeutic approaches decreased p-ERK in SW837 xenograft tumors by 4 hours post last-dose (mean of control: 109, BI-3406 83.5, adagrasib: 69.74, cetuximab: 58.19, BI-3406 plus adagrasib: 66.74 and cetuximab plus adagrasib: 50.8) (**Extended Data Fig 3b)**, a finding consistent with that found in the F3008 CRC PDX model using a multiplex immunofluorescence (mIF) assay (**Fig 4f**).

Although the combination approaches induce enhanced anti-proliferative effects, re-activation of MAPK pathway signaling, a mechanism associated with therapeutic resistance to KRAS^G12C^ targeted therapy^19,21^, was observed in both NCI-H2122 and SW837 xenograft models. In NCI-H2122 xenograft tumors, a slight rebound of MAPK pathway activity was observed at 48 hours after the last dose of adagrasib plus BI-3406 (median MPAS=−0.21) or TNO155 (median MPAS= −0.12) (**Fig 4b,c**), which was consistent with *in vitro* findings (**Fig 3b**). In SW837 xenograft tumors, a modest rebound in MAPK pathway activity, as well as upregulation of SOS1/2 and RTKs (e.g. ERBB2 and ERBB3), were observed by 48 hours after the last dose of adagrasib alone or with BI-3406 or cetuximab (**Extended Data Fig 1b,c**). Additionally, a partial rebound of p-ERK levels was observed at 48 hours after the last dose of adagrasib alone or with BI-3406 in SW837 xenograft models (**Extended Data Fig 3b)**. Combination approaches resulted in the increased expression of a range of genes (325 genes in vehicle vs adagrasib plus BI-3406; 330 genes vehicle vs adagrasib plus TNO155) that included genes associated with the MAP kinase family, tumor growth, and cancer progression^32–35^ in both models (**Fig 4c, Supplementary Table 2, Supplementary Table 3**). The emergence of transcriptomic changes associated with therapeutic resistance to KRAS^G12C^ targeted therapy in SW837 models treated with combination approaches is especially interesting, as outgrowth in these tumor-bearing animals was not observed in response to combination treatments within the 42-day experimental period (**Fig 2b, Supplemental Table 3**).

Overall, our findings underscore the necessity of co-administering adagrasib with inhibitors targeting regulators of RAS GTP-loading to improve the efficacy and durability of KRAS^G12C^ targeted therapy. When compared to single-agent approaches, combination treatment induced more profound gene modulation *in vivo* accompanied by strengthened inhibition on the RAS-MAPK pathway, leading to enhanced anti-tumor efficacy. Upregulation of MAPK pathway signaling, as well as pharmacodynamic markers associated with MAP kinase family and cancer progression, were observed in response to all treatment types. Additional dedicated studies are needed to better understand the mechanistic connection between these transcriptomic changes and the emergence of outgrowth in KRAS^G12C^-mutated models treated with combination therapies.

### KRAS^G12C^i-resistant cells respond to KRAS^G12C^i plus SOS1i combination therapy

As rapid resistance to KRAS^G12C^i monotherapy was observed in NSCLC and CRC models *in vitro* (**Fig 1d; Fig 3**) and *in vivo* **(Fig 2; Fig 4b,c; Extended Data Fig 1b,c; Extended Data Fig 3b**), we sought to investigate whether combination therapy with SOS1i could be leveraged to induce anti-tumor effects in tumors with acquired KRAS^G12C^i resistance.

We first evaluated the effectiveness of combination therapy in preventing cellular outgrowth due to the acquisition of secondary KRAS mutations that have been shown to drive resistance to adagrasib and sotorasib^13,14,36^. To this end, we constructed KRAS^G12C^ Ba/F3 transgenic cell pools that harbored a comprehensive set of secondary KRAS mutations, and assessed the response of the KRAS^G12C^ Ba/F3 clone library to KRAS^G12C^i (adagrasib or sotorasib) alone or combined with escalating doses of BI-3406. Consistently, when compared to KRAS^G12C^i alone, combination therapy resulted in a strong reduction in the number of outgrowth-positive wells that had been seeded with random library sub-pools in a dose-dependent manner, with KRAS^G12C^i plus 600 nM BI-3406 producing stronger inhibition than 300 nM BI-3406 (**Extended Data Fig 4a**). These findings demonstrate the benefit of combining BI-3406 with a KRAS^G12C^i to treat *KRAS^G12C^*-mutated tumors that have acquired resistance to KRAS^G12C^i monotherapy due to the acquisition of secondary KRAS mutations that may impede the ability of the inhibitor to bind and inhibit KRAS.

Subsequent studies evaluated the potential of leveraging combination therapy to overcome acquired resistance to KRAS^G12C^i. Adagrasib-resistant NCI-H358 cells were generated by long-term culture in high-dose adagrasib (10x IC_50_) and their response to mono- or combination therapy were then assessed. All remaining NCI-H358 cells following long-term culture retained KRAS^G12C^ and, as expected, displayed less sensitivity to adagrasib than parental NCI-H358 cells (**Extended Data Fig 4b**). Adagrasib-resistant NCI-H358 cells were more sensitive to adagrasib plus BI-3406 treatment compared to adagrasib alone, although this sensitivity was not as strong as the response of parental (adagrasib naïve) cells to KRAS^G12C^i monotherapy (**Extended Data Fig 4b)**. Similar results were obtained from adagrasib-resistant NCI-H358 cells treated with adagrasib plus TNO155 (**Extended Data Fig 4b)**. Together, these data suggested that the profound suppression of KRAS signaling induced by adagrasib plus BI-3406 combination treatment may be effective in KRAS^G12C^-mutated NSCLC with acquired adagrasib resistance.

We next evaluated whether combination with BI-3406 was effective in re-establishing sensitivity to adagrasib in tumors that gained acquired resistance to adagrasib *in vivo*. SW837 xenograft tumors were treated with adagrasib, resulting in an initial phase of tumor control followed by outgrowth. Regrowth of more than 100mm^3^ was considered to represent acquired resistance.

Relapsed tumors were then randomized on Day 63 and 84 for retreatment with adagrasib alone or in combination with either BI-3406, cetuximab, or TNO155 (**Fig 5a, Extended Data Fig 4c**). Treatment of adagrasib plus BI-3406 or TNO155, but not adagrasib alone, resulted in the regression or relapsed tumors **(Fig 5b; Extended Data Fig 4d)**, whereas tumor stasis was observed in animals treated with adagrasib plus cetuximab **(Fig 5b)**. These findings suggest that combinations with BI-3406 or TNO155 can be effective at re-establishing tumor growth control in tumors with acquired resistance to adagrasib.

**Figure 5.**
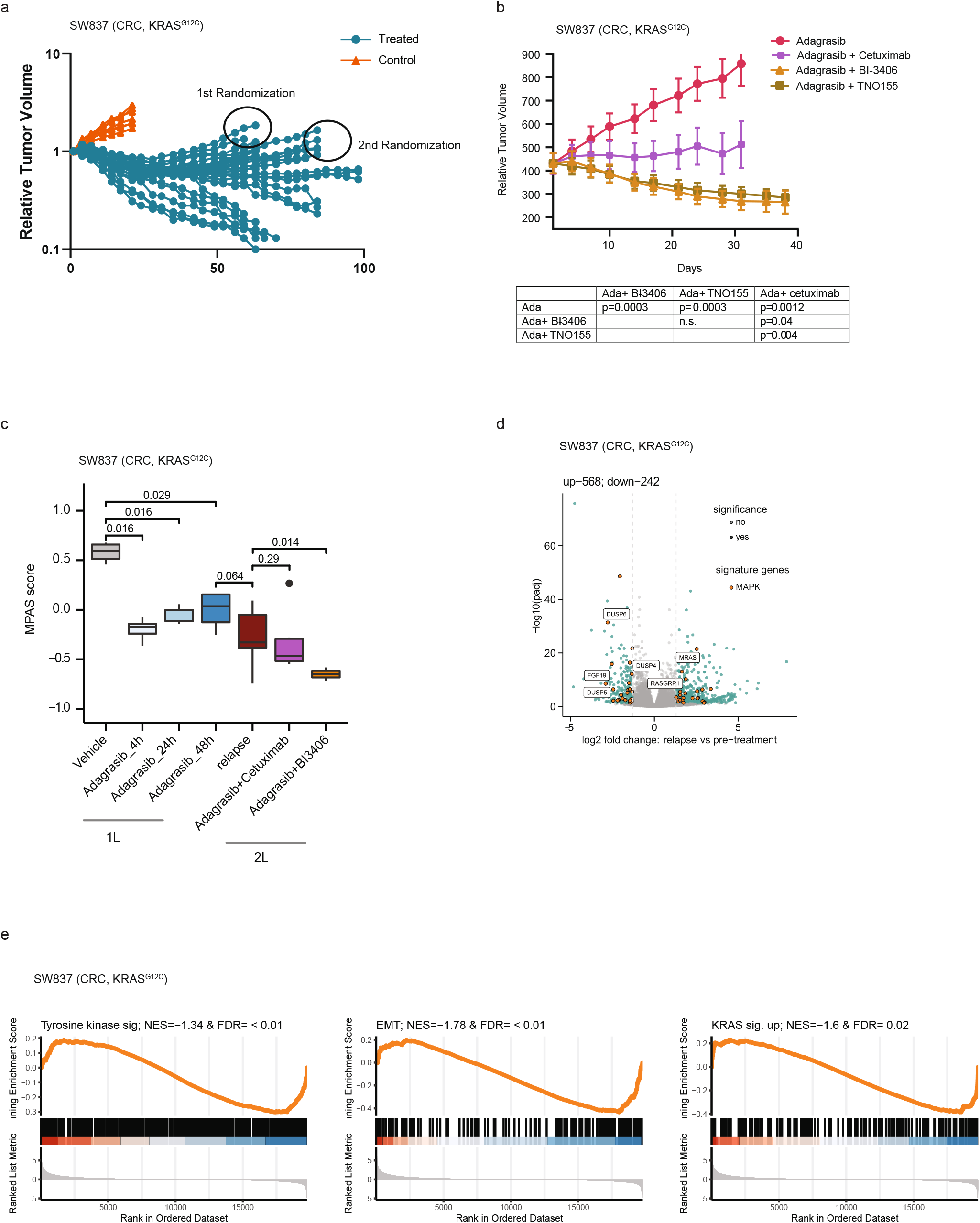
Co-administration of BI-3406 with adagrasib leads to regressions in KRAS^G12C^i-resistant colorectal cancer (CRC) models. **a**, Tumor growth into the KRAS^G12C^-mutant CRC xenograft model (SW847) treated with vehicle (red) or adagrasib (blue; 50 mg/kg, 5 days on/2days off per week). N=30 per group (n=10). **b**, Outgrowing adagrasib-resistant tumors from (a) that had shown an increase in size at least of 100mm^3^, were treated with adagrasib (50 mg/kg, qd, p.o.), adagrasib (50 mg/kg, qd, p.o.) plus BI-3406 (50mg/kg, bid, p.o.), adagrasib (50 mg/kg, qd., p.o.) plus cetuximab (20mg/kg, twice weekly, i.p.), or adagrasib (50 mg/kg, qd) plus TNO155 (10mg/kg, bid). N=5 tumors/group. Mean ± SEM are shown. Data were analyzed on Day 31 by an one-sided non-parametric Mann-Whitney-Wilcoxon U-tests adjusted for multiple comparisons according to the Bonferroni-Holm Method within each subtopic; ada = adagrasib; n.s. = non-significant. **c**, Modulation of overall MAPK pathway activity, as indicated by the MAPK Pathway Activity Score (MPAS), in tumors derived from SW837 xenograft models. Tumors were collected from models before adagrasib treatment (pre-treatment) and during relapse as well as from adagrasib-resistant models treated with adagrasib plus BI-3406 or cetuximab. N=3-16/group. Boxplots show low and upper quartiles and median line is indicates. Whiskers: 1.5 x interquartile range. Data analyzed by two-sided Wilcoxon ran sum test. **d**, Volcano plot of differentially regulated genes between pre-adagrasib treatment (pre-treatment) and relapsed SW837 tumors. The most significantly differential genes involved in MAPK pathway are highlighted. **e**, Gene set enrichment analysis (GSEA) plots show downregulated pathways between tumors collected from SW837 models prior to adagrasib treatment and during relapse after chronic adagrasib treatment. EMT = epithelial-mesenchymal transition. NES = normalized enrichment score; FDR = false discovery rate-adjusted q value.

The molecular mechanism underlying the strong anti-proliferative response observed in relapsed SW837 xenograft models upon KRAS^G12C^i plus SOS1i combination therapy was then investigated. Initially, we sought to delineate the differential transcriptomic profiles underlying acquired resistance to adagrasib monotherapy in our models. In this setting, we failed to identify secondary KRAS mutations or enriched KRAS expression in relapsed tumors, events previously reported in tumors with acquired resistance to KRAS^G12C^i^13^ (**Supplementary Table 4**). Interestingly, MAPK pathway activity in adagrasib-resistant relapsed SW837 xenograft tumors was significantly lower than that observed in vehicle-treated tumors, but in a similar range to that in parental SW837 xenograft tumors at 48 hours post-last dose (**Fig 5c)**. Together with our previous findings (**Extended Data Fig 1b,c**), these data indicate that the modest rebound in overall MAPK pathway activity in parental SW837 xenograft tumors at 48 hours after the last dose of adagrasib did not further increase during the acquisition of KRAS^G12C^i resistance in relapsed tumors. Administration of adagrasib plus BI-3406 (p=0.014), but not cetuximab (p=0.29), to SW837 xenograft models further downregulated MAPK pathway activity when compared to levels in relapsed models (**Fig 5c**). These data are consistent with results showing that co-administration of BI-3406 with adagrasib was capable of inducing tumor regression in relapsed SW837 xenograft tumors (**Fig 5b**).

We next sought to gain a more granular understanding of the transcriptomic changes that occur in SW837 tumors relapsing under adagrasib treatment. Identification of significantly enriched biological pathways was conducted by leveraging gene set enrichment analysis (GSEA) in a supervised manner on gene sets associated with KRASG12Ci^13,15,37,38^. Our findings revealed that gene sets associated with tyrosine kinase signaling (n=543), EMT (n=204), as well as genes known to be regulated by active KRAS (n=220) were downregulated in relapsed tumors, but no specific pattern of downregulation could be discerned (**Fig 5e**). Differential expression analysis revealed that 810 genes were significantly deregulated (242 and 568 genes down and upregulated, respectively) in tumors undergoing relapse (**Fig 5d, Supp. Table 5**). Interestingly, we found that DUSP5 and DUSP6, two pharmacodynamic biomarkers of KRAS^G12C^i, were downregulated in relapsed tumors (**Fig 5d; Extended Data Fig 4e**), which, together with the fact that no secondary KRAS mutations were detected, suggests that adagrasib still binds and inhibits KRAS^G12C^. Notably, upregulated genes in relapsed SW837 xenograft tumor included MRAS, a homolog of the RAS family of GTPases^39^, and RasGRP1, a Ras guanine nucleotide exchange factor (GEF)^40,41^, but not NRAS nor HRAS (**Fig 5d; Extended Data 4e; Supplementary Table 5**). Together, these data outline the complex ways that KRAS^G12C^i and its combination partners regulate members of the MAPK pathway in relapsed SW837 xenograft tumors.

In summary, we demonstrate that adagrasib plus BI-3406 or TNO155, but not adagrasib alone or a combination with cetuximab, results in enhanced inhibition on the MAPK pathway and subsequently induced tumor regression in a SW837 xenograft model with acquired resistance to adagrasib. Interestingly, our data indicate that the acquired adagrasib resistance in our models is likely not mediated by the gain of secondary KRAS mutations nor by a strong rebound in MAPK signaling. Additional investigation is necessary to clarify the contribution of specific genes towards the development of KRAS^G12C^i resistance and relapse.

## Discussion

Efforts to target mutant KRAS signaling in cancer have advanced tremendously in the past few years, with multiple KRAS^G12C^-specific inhibitors currently in clinical or preclinical development^5,9,11,42–45^. In 2^nd^ line patients with KRAS^G12C^-mutated NSCLC, adagrasib (MRTX849) and sotorasib (AMG 510) have achieved objective response rates of 45% and 37% and disease control rates of 96% and 81%, respectively^10,11^, but efforts are needed to improve the rate and duration of response to KRAS^G12C^-specific inhibitors. Here, we show that co-administration of BI-3406, a SOS1 inhibitor, with adagrasib in preclinical KRAS^G12C^ mutant NSCLC and CRC models enhances the potency and duration of the anti-tumor response to levels better than that induced by adagrasib alone, and comparable to that induced by combination approaches (e.g. KRAS^G12C^i plus SHP2i or EGFRi) that are currently evaluated in the clinic^21,25,26^.

Our findings indicate that the enhanced anti-tumor effects induced by co-administering BI-3406 with adagrasib in NSCLC and CRC models are driven by enhanced suppression of RAS-GTP levels and subsequent downstream signaling. Compared to that induced by single-agent treatments, concurrent inhibition of SOS1 and KRAS^G12C^ further reduced RAS-GTP levels and additionally downregulated pharmacodynamic biomarkers of the MAPK pathway and mitosis (e.g. p-ERK, ETV4, ETV5, EGR1, and Ki-67). These findings are consistent with previous studies where mutant KRAS^G12C^ inhibitors were combined with partners that target SHP2 or RTKs^5,19,21^, and suggest that KRAS^G12C^i plus SOS1i combination treatment may lead to a more profound inhibition of KRAS downstream signaling, thereby potentially eliciting stronger and more durable responses in the clinic. Our promising preclinical findings provide support to the progression of BI 1701963, a clinical stage SOS1 inhibitor, in combination with sotorasib (NCT04185883) or the KRAS^G12C^i BI 1823911 (NCT04973163) into clinical testing in patients with advanced solid tumors harboring a KRAS^G12C^ mutation.

Early clinical data^5,13^ and preclinical studies^14,^*15* demonstrating the rapid onset of acquired resistance to KRAS^G12C^i monotherapy as well as the limited duration of progression-free survival of patients treated with sotorasib^9,11^ underscore the need to identify rational combination strategies that can extend and improve response to KRAS^G12C^i targeted therapy. The enhanced and extended anti-tumor response induced by combining KRAS^G12C^i with BI-3406 or TNO155, was most pronounced in our CRC model. Impressively, BI-3406 plus adagrasib combination treatment extended the anti-proliferative response to KRAS^G12C^ inhibition in our CRC model, as tumor regression in animals treated with SOS1i plus adagrasib combination therapy was still evident at time points when relapse was observed in those treated with adagrasib alone. The ability of a SOS1i or SHP2i to regulate downstream signaling from multiple RTKs may underly the enhanced anti-tumor response observed^25,46^. While our findings are consistent with studies assessing the anti-tumor responses of KRAS^G12C^i combined with agents that inhibit SHP2, EGFR, MEK, PI3K, or CDK4/6^19,21,22,47^, inhibiting SOS1, which directly regulates GTP loading onto KRAS, may avoid the pleiotropic effects of SHP2 inhibitors^48,49^, which may result in better tolerance in the clinic. Altogether, our study positions SOS1i plus KRAS^G12C^i combination treatment as an effective strategy for extending the anti-tumor response in patients harboring KRAS^G12C^-mutant CRC.

Acquired resistance to KRAS^G12C^i monotherapy is thought to be mediated by multiple mechanisms^5,11,13,14,19–21^, such as upstream bypass events including RTK activation^21^, KRAS^G12C^ amplification^50^, activation of alternative signaling pathways including PI3K, mTOR, YAP-TEAD pathways, acquisition of low frequency secondary KRAS mutations that preclude compound binding (e.g. R68S, Y96C/D, H95D/Q/R)^13^, conversion of the G12C codon mutation, and additional activating KRAS mutations (G13D, Q61H)^15,36^. None of these mechanisms emerged as the primary mechanism of resistance in our models. In an attempt to establish the ability of SOS1i to address some of these mechanisms, we demonstrate here that co-treatment with a SOS1i could provide benefit to patients who progress on KRAS^G12C^i due to acquired secondary on-target mutations in the KRAS^G12C^ setting. Co-administration of BI-3406 with a KRAS^G12C^i further inhibited the outgrowth of KRAS^G12C^-driven cells with randomly generated secondary mutations, likely due to its ability to enhance KRAS blockade in double-mutants that may impair the binding of KRAS^G12C^ inhibitors.

A moderate but rapid rebound of pharmacodynamic markers associated with MAPK pathway signaling, parallel MAPK pathways, and cancer progression emerged as a probable contributor to acquired resistance to adagrasib in our CRC models. This hypothesis is supported by evidence of upregulated RasGRP1 and MRAS expression in relapsed SW837 xenograft tumors accompanying a modest rebound in MAPK pathway activation. Although possessing GEF activity similar to SOS1, RasGRP1 has been shown to suppress EGFR-SOS1-RAS signaling in normal intestinal epithelial and CRC cells^41,51^, and reported to limit CRC cell proliferation *in vitro* and CRC xenograft tumor growth*^41^*. Instead, RasGRP1 can directly bind to and facilitate the GTP-loading of MRAS, a homolog of the RAS family of GTPases^39,40^. MRAS, together with SHOC2 and PP1C, forms a phosphatase complex that contributes to the increased activation of RAS-MAPK signaling through dephosphorylation of inhibitory sites on RAF kinases^52^. Aberrant activity of this complex has been implicated in Noonan syndrome; a developmental disorder caused by overactive RAS-MAPK signaling^53^. While aberrant MRAS activity in cancer has been rarely documented^39^, low allele frequency mutations in MRAS have been reported to emerge during the development of KRAS^G12C^i resistance in NSCLC cell lines^15^. Thus, the elevated MRAS expression, along with the modest rebound in MAPK activity, observed in our CRC models with acquired adagrasib resistance suggest that MRAS may compensate for loss of KRAS activity and drive re-activation of MAPK signaling, whose activity may still be subjective to the regulation of SOS1^40^. Therefore, while additional functional studies are needed to clarify the role of elevated MRAS, RasGRP1, and RAS-MAPK pathway activity during acquired resistance to KRAS^G12C^i, our findings suggest that combination with SOS1i is an attractive combination strategy to potentially overcome acquired resistance to KRAS^G12C^i.

In summary, we demonstrate that co-administering BI-3406 with adagrasib elicits an enhanced anti-tumor response in both adagrasib-responsive and -resistant KRAS^G12C^-driven tumors.

Moreover, response to adagrasib plus BI-3406 is comparable to that of adagrasib with TNO155 and stronger than that observed in response to adagrasib and cetuximab, which has elicited impressive response in KRAS^G12C^-driven CRC patients^26^. Together, our findings underscore the potential of leveraging a SOS1i combination partner as a strategy to increase the efficacy and duration of KRAS^G12C^ targeted therapy in patients harboring KRAS^G12C^-driven NSCLC or CRC. Follow-up studies are needed to assess the potential of combining SOS1 inhibitors with “KRAS^G12C^-on” inhibitors, a new class of KRAS allele-specific inhibitors that target the active GTP-bound state^54^. Early clinical data indicate that co-treatment of KRAS^G12C^-on inhibitors with SHP2i prolongs the durability of response^55^; we hypothesize that combination with an SOS1i may achieve similar results, especially in the setting of tumors harboring secondary KRAS mutations^54^. Overall, we envision that co-administering an SOS1i with allele-specific KRAS inhibitors may improve and prolong clinical responses in patients with KRAS-mutated tumors.

## Acknowledgements

The authors thank R. Mullinax, A. Zuniga, A. Harris, and G. F. Draetta for their support. This study was supported by Boehringer Ingelheim and the Austrian Research Promotion Agency (FFG) with support awards 854341, 861507, 867897, 874517 883626 and 892584. This study was also supported by a sponsored research Alliance collaboration between Boehringer Ingelheim and MD Anderson Cancer Center, with funding provided by Boehringer Ingelheim. This study was also supported by the MDACC Science Park NGS Core grant: CPRIT Core Facility Support award (RP170002) and the RPPA Core facility were funded by NCI #CA16672.

## Author Contributions

M. H. Hofmann, V. Thatikonda, S. Jurado, I. Waizenegger, D. Gerlach, F. Trapani, P. Jaeger, H. Lu, C. P. Vellano, T. P. Heffernan, M.P. Petronczki and J. R. Marszalek contributed to project conceptualization, supervision, and administration. V. Thatikonda, D. Gerlach, M. Gmachl, K. Bosch, S. Lieb, A. Jeschko, P. A, Jaeger, S. Strauss, F. Trapani, H. Lu, N. Feng, S. Kopetz, A. A. Machado, E. D. Marszalek, M. Mahendra, M.P. Petronczki and A. Sorokin contributed to executing methodology and data acquisition. M. H. Hofmann, V. Thatikonda, S. Jurado, K. Kostyrko, I. Waizenegger, K. Bosch, D. Gerlach, F. Trapani, P. Jaeger, H. Lu, C. A. Bristow, and C. P. Vellano contributed to data analysis. V. Thatikonda, S. Jurado, D. Gerlach, M. H. Hofmann, K. Kostyrko I. Waizenegger, P. Jaeger, M. Gmachl, N. Kraut, M. P. Petronczki, M. Pearson, H. Lu, and S. Gao contributed to manuscript and figure preparation. All co-authors reviewed the manuscript prior to submission.

## Competing Interests

V. Thatikonda, S. Jurado, K. Kostyrko, K. Bosch, D. Gerlach, M. Gmachl, S. Lieb, A. Jeschko, P. A. Jaeger, S. Strauss, F. Trapani, M. Pearson, I. Waizenegger, M. P. Petronczki, N. Kraut and M. H. Hofmann report grants from the Austrian Research Promotion Agency (FFG), receive personal fees from Boehringer Ingelheim (full-time employee) during the conduct of the study. M.H. Hofmann and M. Gmachl have been listed as inventor on patent applications for SOS1 inhibitors. A. Sorokin, S. Kopetz, H. Lu, A. A. Machado, M. Mahendra, E. D. Marszalek, S. Gao, N. Feng, C. A. Bristow, C. P. Vellano, T. P. Heffernan, and J. R. Marszalek report other from Boehringer Ingelheim (sponsored research) during the conduct of the study and this work was performed under a sponsored research collaboration between MD Anderson and Boehringer Ingelheim, for which the latter provided funding support. S. Kopetz has ownership interest in Lutris, Iylon, Frontier Medicines, Xilis, Navire and is a consultant for Genentech, EMD Serono, Merck, Holy Stone Healthcare, Novartis, Lilly, Boehringer Ingelheim, AstraZeneca/MedImmune, Bayer Health, Redx Pharma, Ipsen, HalioDx, Lutris, Jacobio, Pfizer, Repare Therapeutics, Inivata, GlaxoSmithKline, Jazz Pharmaceuticals, Iylon, Xilis, Abbvie, Amal Therapeutics, Gilead Sciences, Mirati Therapeutics, Flame Biosciences, Servier, Carina Biotech, Bicara Therapeutics, Endeavor BioMedicines, Numab, Johnson & Johnson/Janssen, Genomic Health, Frontier Medicines, Replimune, Taiho Pharmaceutical, Cardiff Oncology, Ono Pharmaceutical, Bristol-Myers Squibb-Medarex, Amgen, Tempus, Foundation Medicine, Harbinger Oncology, Inc, Takeda, CureTeq, Zentalis, Black Stone Therapeutics, NeoGenomics Laboratories, Accademia Nazionale Di Medicina, and receive research funding from Sanofi, Biocartis, Guardant Health, Array BioPharma, Genentech/Roche, EMD Serono, MedImmune, Novartis, Amgen, Lilly, Daiichi Sankyo. T. P. Heffernan receives advisory fees from Cullgen Inc. and Roivant Discovery.

## Data Availability Statement

Cell line derived xenograft RNA-seq data analyzed in this study deposited to GEO under the id GSEXXXXX. All the computational analysis was performed using R statistical programming language (v4.0.2) with open source packages described the Methods. All other data are available from the corresponding author upon reasonable request.

## Methods

### Cell Lines

Cell lines were purchased from American Type Culture Collection (ATCC). SW837 cells were cultured in DMEM (Sigma #D6429) with 10% FBS (Hyclone, Thermo Fisher Scientific #SH30071.03, Lot #AYS170727). NCI-H2122 cells were cultured in RPMI (Gibco #3A10491-01) + 10% FBS at 37°C, 5% CO_2_ in a humidified incubator. SW837 cells were grown in Leibovitz’s L-15 medium (Gibco #11415-049) for *in vitro* combination studies as shown in Figure 1b (without CO2), for all other studies SW837 cells were grown in DMEM + 10% FCS at 37°C and 5% CO2.

### Western Blot

SW837 or NCI-H2122 cells were seeded in 6 well plates (Corning #3506) at a density of 2.5×10e6 or 1×10e6 cells/well in complete culture medium. After overnight incubation at 37°C and 5% CO_2_ in a humidified incubator, cells were treated with indicated concentrations of MRTX849, BI-3406, TNO155, or SHP099 alone or in combination; control cells were treated with 0.1% DMSO (Sigma-Aldrich #41648). After 6 or 24 hours of treatment, cells were washed with ice cold PBS, harvested, and lysed with MSD lysis buffer (Mesoscale Diagnostics #R60TX-2) containing Tris pH 7.5, 150 mM NaCl, 1 mM EDTA, 1 mM EGTA, 1 % Triton X-100, 10 mM NaF, completed with protease and phosphatase inhibitors (Thermo Fisher Scientific #78440). Protein concentrations were determined via Bradford assay according to manufacturer’s instruction (Bradford Dye, Biorad #5000205). Unless otherwise stated, 20 μg of total protein was separated on a 4-12% polyacrylamide gel (BioRad #3450124) in MOPS Running Buffer (BioRad #1610788) and blotted on a PVDF membrane (BioRad #1704157) with the Bio-Rad Trans-Blot^®^ TurboTM Instrument. For separation of KRAS shift, total protein was separated on a 10% Tris-HCL polyacrylamide gel (BioRad #3450021) in Tris Gycin Running Buffer (BioRad #1610732). Membranes were blocked for 1 hr in 4% skim milk (Millipore #70166) in 1×TBS (BioRad #1706435)/0.1% Tween 20 (Bio-Rad #161-0781) at room temperature and then probed overnight at 4°C with primary antibodies against KRAS (LSBio #LS-C175665; 1:500), pERK Thr202/Tyr204 (Cell Signaling #4376; 1:500), ERK (Cell Signaling #9102; 1000-1:2000), phospho-S6 Ribosomal Protein (Ser235/236) (Cell Signaling #2211; 1:500-1:1000), DUSP-6 (Abcam #ab76310; 1:1000), cleaved PARP (Asp214) (Cell Signaling #9541; 1:1000), Cyclin D1 (Biosite #ARB-Q4OL25-0,5; 1:100); p27 (BD #610241; 1:1000), and ß-Actin (abcam #ab8226; 1:10000). The phosopho-S6 ribosomal protein dilution was prepared in 5% BSA in 1×TBS/0.1% Tween 20, and all other antibody dilutions were prepared in 4% skim milk in 1×TBS/0.1% Tween 20. After washing membranes were then incubated with the following secondary antibodies diluted in respective incubation buffers: goat a-rabbit IgG, HRP conjugated (Dako #P0447; 1:1000), goat a-mouse IgG, HRP conjugated (Dako #P0448; 1:1000). Proteins were visualized using ECL Western Blotting detection reagent (Amersham #RPN2106) according to the manufacturer’s instructions.

### IncuCyte kinetic cell confluence proliferation assay

SW837 and NCI-H2122 cells were seeded in 96 well plates (Corning #3598) at a density of 4000 cells/well in complete culture medium. After overnight incubation at 37°C and 5% CO_2_ in a humidified incubator, cells were treated with indicated concentrations of MRTX849, BI-3406, TNO155, or SHP099 as monotherapy or in combination. Control cells were treated with 0.1% DMSO. Immediately before treatment, cells were stained with IncuCyte^®^ Caspase-3/7 Green Apoptosis Assay Reagent (Essen BioScience #4440) for apoptosis detection according to the manufacturer’s instruction. Cell growth was monitored using the IncuCyte^®^ S3 live cell imaging system (Essen BioScience). Using the 10× objective, two regions of view were collected per well every 4 hours for at least 7 days, with an extended collection period in cases of slow cell growth. Phase contrast and green channel (Ex: 440/80 nm; Em: 504/44 nm) were collected for each experiment. Data were analyzed using Incucyte 2019B software. Values from both regions of each well were averaged and confluence was calculated as the percentage of the image area that was occupied by objects (Phase area confluence). Apoptotic events were calculated as the percentage of the image area that was occupied by green objects.

### Combination Studies – Cell Culture

H2122 cells (human lung cells, ATCC CRL-5985) were grown in RPMI-1640 medium (Gibco #A10491) with 10% FCS (Hyclone #SH30084.03n) at 37°C with 5% CO_2_. After expansion and passaging twice a week at 1:2-1:4, we seeded 800 cells/well in 60 μl in 384-well plates (Greiner #781182) for the proliferation assay. After 24 hours, compounds were added with an ultrasonic dispersion system (Echo, Labcyte System) and incubated for another 72 hours. Cells were then stained with CellTiter-Glo Reagent according to the manufacturer’s protocol (Promega, #G924C) and incubated 15 minutes under shaking. Plates were then read with a plate reader (EnVision, PerkinElmer HTS Multilabel). SW837 (human colon cells, ATCC CCL-235) were grown in Leibovitz’s L-15 medium (Gibco #11415-049) with 10% FCS (Hyclone #SH30084.03n) at 37°C without CO_2_ for combination studies as shown in Figure 1b, for all other studies SW837 cells were grown in DMEM + 10% FCS at 37°C and 5% CO_2_. After expansion and passaging twice a week at 1:2-1:3, we seeded 4000 cells/well in 180μl in 96-well plates (Greiner #655090) for the proliferation assay. After 24 hours, compounds were added with an ultrasonic dispersion system (Echo, Labcyte System) and incubated for another 96 hours. Cells were then stained with PrestoBlue Reagent according to manufacturer’s protocol (InVitrogen cat #A13262) and incubated 4-5 hour in incubator at 37°C, before being read with a plate reader (EnVision, PerkinElmer HTS Multilabel).

### Growth Curve Fitting and Combination Scoring

Curves were fitted using standard nonlinear regression with a four-parameter fit after normalization to percentage of maximum signal (the negative DMSO controls) for each treatment type. The combination cScore quantifies the deviation of the observed effect of two drugs combined when compared to the expected effect based on the observed monotherapies at the same concentration. For this, the measured cell viability is subtracted from the expected cell viability using the Bliss Independence Model^56^, creating a “gap” matrix with the two monotherapy concentrations on the x- and y-axis respectively and the unit [%PoC]. The cScore is then defined as the average gap value of a 3×3 field matrix around the IC_50_ of both compounds, or, if one or both compounds have no measurable IC_50_, the scores at the highest two concentrations of the Gap table. If the IC_50_ is close to the C_max_ or C_min_, the field used for averaging may be 2×3 or 2×2, but never less than 4. This approach focuses on drug combination effects around the IC_50_ values of the respective monotherapies. Positive cScores thus express the average increased potency of the combination over the monotherapies around the IC_50_ in % cell viability PoC.

### Oncogenic KRAS variant generation and viability

Ba/F3 cells (DSMZ, cat. #ACC300) were grown in RPMI-1640 medium supplemented with 10% FCS at 37 °C in 5% CO_2_ atmosphere in the presence of 10 ng/mL IL-3 (R&D). Ba/F3 cells were transduced with retroviruses encoding pMSCV_KRAS_G12C_single site variant Library (ssvL) cloned at TWIST Biosciences and harboring a pool of each possible variant at any AA position of the KRAS gene, a puromycin resistance gene, and the green fluorescent protein (GFP). Platinum-E cells (Cell Biolabs) were used for retrovirus packaging. Retrovirus and 4 μg/mL polybrene were added to Ba/F3 cells for spinfection. Infection efficiency was confirmed by measuring GFP-positive cells using a cell analyzer. Cells with a library representation of >1000x were further cultivated in the presence of puromycin (1 μg/mL) to select for transduced cells. Following selection, IL-3 was withdrawn from transgenic Ba/F3 cells expressing the oncogenic KRAS variants (KRAS ssvL Ba/F3) to make cells dependent on transgene activity. For colony growth assays, KRAS ssvL BaF/3 cells were seeded into 5x 96-well plates at 250 cells/100 μL in growth media per condition. Compounds were added alone or in combination with two different doses of SOS1i. Treated cells were incubated for 14 days at 37 °C with 5% CO_2_ with an addition of 100 μl medium/cpd mixture after 7 days. AlamarBlue™ Cell Viability assay (ThermoFisher) was performed, and after 6 hours, fluorescence was measured by using the multilabel Plate Reader VICTOR X4. The raw data were imported into Microsoft Excel and the signal to background ratio was calculated. Ratios >1.5 were counted as colony growth, and then visualized in PRISM, GraphPad Inc.

### Cell line-derived efficacy studies and biomarker studies in mice

Female BomTac:NMRI-*^Foxn1nu^* mice were used for all xenograft studies. For biomarker and efficacy experiments using SW837 or NCI-H2122 tumor-bearing mice, female mice were engrafted subcutaneously with 5 million cells suspended in Matrigel (Corning, #356231) diluted in 1xPBS with 5% FCS. Tumors were randomized by tumor size in groups using the automated data storage system Sepia. Mice were treated once at time point 0 hour (qd) and in addition 6 hours later in case of twice daily treatment. Tumor size was measured by an electronic caliper and body weight was monitored daily. The analysis follows largely the procedures described previously^57,58^. BI-3406, adagrasib, and TNO155 were dissolved in 0.5% Natrosol. BI-3406 was administered at 50 mg/kg p.o. twice a day, adagrasib at 100 mg/kg p.o. once per day, and TNO15 at 10 mg/kg, bid. Trametinib was dissolved in 0.5% DMSO and 0.5% Natrosol and dosed twice daily with 0.1 mg/kg. Cetuximab was dissolved in 0.9% NaCl and dosed at 15 mg/kg i.p. twice a week. The control group was treated with 0.5% of Natrosol orally in the same frequency as in the treatment groups (twice daily). All compounds were administered either intragastrically by gavage (10 mL/kg) or intraperitoneally (Cetuximab). Mice included in the biomarker studies were treated for 7 continuous days. Tumors were explanted at 4h, 24h, or 48 hours post last dose, and were either embedded in paraffin or fresh frozen for further analysis.

For investigating the efficacy of treatments on SW837 xenograft models with acquired adagrasib resistance, animals bearing established SW837 tumors were treated long-term with 50 mg/kg adagrasib for 5 days on/2days off per week. Tumor size was measured by an electronic caliper and body weight was monitored daily. Most tumors first underwent regression, with outgrowth occurring after several weeks of treatment. Outgrowing tumors that reached an increase of tumor size of at least 100 mm^3^ compared to the smallest size of the respective tumor then were randomized on day 63 and 84 for inclusion in treatment groups for a 2^nd^ treatment. Results of the efficacy experiments starting on day 63 and day 84 were combined for analysis as no difference was observed in the outcome. All animal studies were approved by the internal ethics committee and the local Austrian governmental committee. Sample sizes were determined by performing power analysis.

### PDX Studies

PDX model characterization and profiling have been described previously ^27^. F3008 and B8032 PDX tumor fragments (4 × 4 × 4 mm^3^) were implanted on the right hind flanks of NSG female mice (Jackson Laboratory) and allowed to grow to an average volume of 100–250 mm^3^ as monitored by caliper measurements. All P.O. compounds were dosed on a 5 days on/2 days off schedule. At enrollment, animals were randomized and treated with vehicle (0.5% Natrosol) P.O. twice a day (6 hours apart), BI-3406 at 50 mg/kg P.O. twice a day (6 hours apart), cetuximab at 15 mg/kg I.P. twice a week, adagrasib at 100 mg/kg P.O. once per day, or adagrasib (P.O. once per day) plus BI-3406 at 50 mg/kg P.O. twice a day (6 hours apart) or cetuximab at 15 mg/kg I.P. twice a week. Mice were 11 weeks old and treatment group sizes included at least 5 to 8 mice per group. All animals received LabDiet 5053 chow ad libitum. Adagrasib was purchased from MedChemExpress and cetuximab was purchased from MD Anderson Cancer Center pharmacy and BI-3406 was synthesized at Boehringer-Ingelheim. During the PDX studies, tumor growth was monitored twice a week with calipers and the tumor volume (TV) was calculated as TV = (D × d^2^/2), where “D” is the largest and “d” is the smallest superficial visible diameter of the tumor mass. All measurements were documented as mm^3^. Body weights were measured twice weekly and used to adjust dosing volume and monitor animal health. All animal studies were approved by the internal ethics committee and the local Austrian governmental committee. Sample sizes were determined based on a previous, similar study^27^.

### Pharmacodynamic (PD) Biomarker Analysis

For the PD biomarker studies using F3008 CRC PDX models, tumor fragments (4 × 4 × 4 mm^3^) were implanted on the right hind flanks of NSG female mice (Jackson Laboratory) and allowed to grow to an average volume of 250–350 mm^3^ as monitored by caliper measurements. At enrollment, animals were randomized and treated for 5 days with vehicle (0.5% Natrosol) P.O. twice a day (6 hours apart), adagrasib at 100 mg/kg P.O. once per day, or adagrasib plus BI-3406 at 50 mg/kg P.O. twice a day (6 hours apart) or cetuximab at 15 mg/kg I.P. on Day 1 and Day 4. Tumors were collected 4 hours after the last dose on the fifth day of treatment and fixed in 10% neutral buffered formalin overnight and then processed and embedded in paraffin. Formalin fixed and paraffin embedded (FFPE) blocks were sectioned into 3 μm thick sections, deparaffinized, then rehydrated by serial passage through xylene and graded alcohol. Sections were subjected to an initial heat-induced epitope retrieval (HIER) in citrate buffer, pH 6, at 95°C for 15 minutes. Anti-phospho-p44/42 MAPK (ERK1/2) (Thr202/Tyr204) (1:2000, Cell Signaling Technology #4370) was developed using Opal tyramide signal amplification (TSA) followed by direct immunofluorescence of HLA conjugated to Alexa 647 (1:250, Abcam #199837). RNAscope in situ hybridization assay was performed following manufacturer’s protocol (Advanced Cell Diagnostics, Inc) using DUSP6 (cat# 405361), EGR1 (cat# 457671-C2), and POLR2A (cat# 310451-C4) probes. Appropriate positive and negative controls were included with the study sections. Digital images of whole-tissue sections were acquired using Vectra Polaris Automated Quantitative Pathology Imaging System (Akoya Biosciences) and representative regions were selected for each whole slide and processed using inForm Software v2.4 (Akoya Biosciences). Processed images were then analyzed using HALO Software v3.2 (Indica Labs Inc.).

For PD biomarker studies using NCI-H2122 and SW837 models, animals were first randomized and treated for 5 days with dosing schedules described above. Tumors were collected at 4, 24, and 48 hours after last dose of treatment. Tumor samples were embedded in FFPE and processed as described above. Briefly, (FFPE) blocks were sectioned into 3 μm thick sections, deparaffinized, and rehydrated by serial passage thorough xylene and graded alcohol. Sections were then subjected to heat induced antigen retrieval and stained for anti-phospho-p44/42 MAPK (ERK1/2) (Thr202/Tyr204) and anti-KI-67 (CST #9027, 1/400 in PBS/2% BSA) on bond RX (LEICA). Appropriate negative and positive controls and isotype controls were included. Digital images of whole tissue sections were acquired with NanoZoomer scanner (Hamamatsu) and processed images were then analyzed using HALO Software v3.2 (Indica Labs Inc.). Results were reviewed by a pathologist.

### RNA isolation and sequencing library preparation for expression profiling

Cell line-derived xenograft samples for expression profiling were prepared with either the QuantSeq or TruSeq protocol. QUANT-seq libraries were prepared as previously described^27^. Briefly, cells were lysed in TRI Lysis Reagent (Qiagen, #79306) according to the manufacturer’s instructions. Instead of chloroform, 10% volume 1-bromo-2-chloropropane (Sigma-Aldrich, #B9673) was added. Total RNA was isolated with RNAeasy Mini Kit (Qiagen, #73404). Quant-seq libraries were prepared using the QuantSeq 3′ mRNA-Seq Library Prep Kit FWD for Illumina from Lexogen (#015.96) according to the manufacturer’s instructions. Samples were subsequently sequenced on an Illumina NextSeq 500 System with a single-end 76 bp protocol. For TruSeq, a similar RNA isolation protocol was used and RNA-seq libraries were prepared using TruSeq RNA library preparation kit v2 according to the manufacturer’s instructions.

### Data Analyses

Statistical analyses were performed with R v4.0.2 and Bioconductor 3.7 or GraphPad Prism (v9.3.1). Associations of gene mutations with the sensitivity status of cell lines as well as comparisons of tumor volumes between control and experimental groups were analyzed with a Fisher exact test. All other data meeting the requirements for parametric analyses were assessed with paired Student’s t-tests or one-way analysis of variance (ANOVA). All statistical analyses used absolute values when calculating tumor volume. Datasets that deviated from normal distribution were analyzed with the non-parametric two-sided Wilcoxon rank sum test. When applicable, p values were adjusted multiple comparisons according to Bonferroni–Holm, Benjamini-Hochberg False Discovery Rate analysis, or Tukey’s multiple comparisons test. The level of significance was fixed at α = 5% such that an (adjusted) p value of less than 0.05 was considered to be statistically significant. Differences were observed to be indicative whenever 0.05 ≤ P < 0.10.

Gene expression (RNA-seq) analysis was performed as previously described^27^. Briefly, reads from grafted samples were filtered into human and mouse reads using Disambiguate. The filtered reads were then processed with a pipeline by building upon the implementation of the ENCODE’ “Long RNA-seq” pipeline; filtered reads were mapped against the Homo sapiens (human) genome hg38/GRCh38 (primary assembly, excluding alternate contigs) or the Mus musculus (mouse) genome mm10/GRCm38 using the STAR (v2.5.2b)^59^ aligner allowing for soft clipping of adapter sequences. For quantification, we used transcript annotation files from Ensembl version 86, which corresponds to GENCODE 25 for human and GENCODE M11 for mouse. Samples were quantified with the above annotations, using RSEM (v1.3.0) and featureCount (v1.5.1)^60^. Quality controls were implemented using FastQC (v0.11.5), picardmetrics (v0.2.4) and dupRadar (v1.0)^61^ at the respective steps.

Differential expression analysis was performed on the human mapped counts derived from featureCounts using DESeq2 (v1.28.1)^62^. We used an absolute log2 fold change cut-off of 1.3 and a false discovery rate (FDR) of <0.05.

Gene set enrichment analysis was performed using fgsea^63^ R/Bioconductor (v1.14) package and hallmark gene sets from the molecular signatures database (MSigDB v7.5.1^64^). The resulting nominal p values were adjusted using Benjamini-Hochberg multiple testing correction method and gene sets with adjusted p value < 0.01 were considered as significant.

Heatmaps of GSEA analysis were generated using ComplexHeatmap (v2.4.3)^65^ and upset plots were generated using UpSetR (v1.4.0)^66^ R/Bioconductor packages. All other figures from data analyses were visualized using ggplot2 (v3.3.2)^67^ R package.

## Extended Data Figure Legends

**Extended Data Figure 1.**
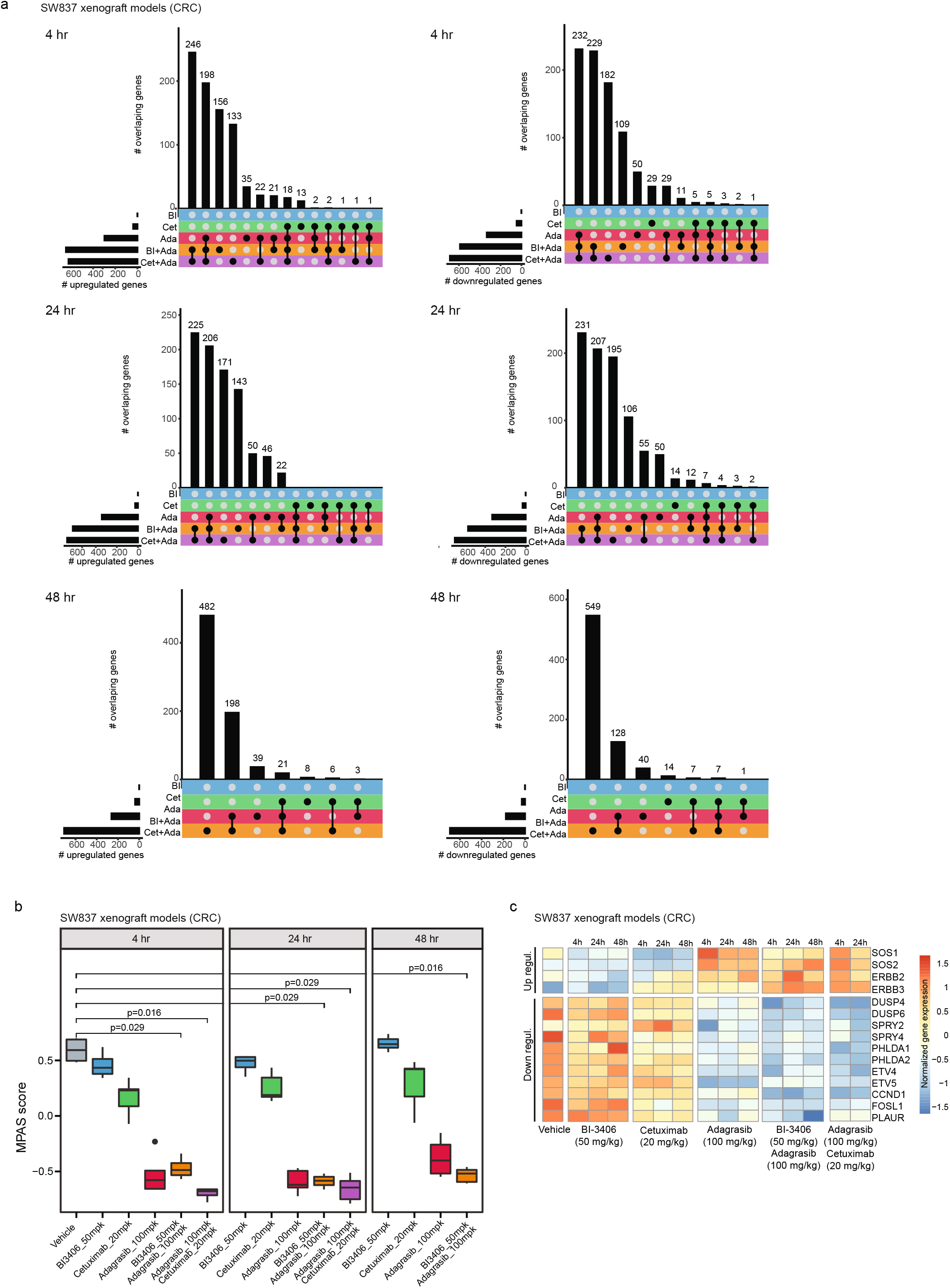
Gene modulation by single-agent or combination treatments in a KRAS^G12C^–driven colorectal cancer (CRC) xenograft model. All treatments in SW837 xenograft models were administered over seven consecutive days. **a**, Overlap of up-regulated (left) or down-regulated (right) genes in tumors from SW837 xenograft models across three time points after receiving one of these treatments: 50 mg/kg bid, p.o. BI-3406 (BI); 20 mg/kg twice a week, i.p. cetuximab (Cet); 100 mg/kg qd, p.o. adagrasib (Ada); 50 mg/kg BI-3406 plus 100 mg/kg adagrasib (BI + Ada); 20 mg/kg cetuximab plus 100 mg/kg adagrasib (Cet + Ada). Samples were collected at 4, 24, and 48 hours after the last dose. n = 4– 5 animals/groups. **b**, Modulation of overall MAPK pathway activity, as indicated by the MAPK Pathway Activity Score (MPAS), in tumors from SW837 xenograft models treated with indicated single-agent or combination treatments. Samples were collected at 4, 24 and 48 hours after the last dose. N=4–5/group. Boxplots show low and upper quartiles and median line is indicates. Whiskers: 1.5 × interquartile range. Data analyzed by two-sided Wilcoxon rank sum test. **c**, Differential modulation of select MAPK pathway genes in tumors from SW837 xenograft models treated with indicated single-agent or combination treatments. Samples were collected at 4, 24, or 48 hours post last dose. N=4–5/group

**Extended Data Figure 2.**
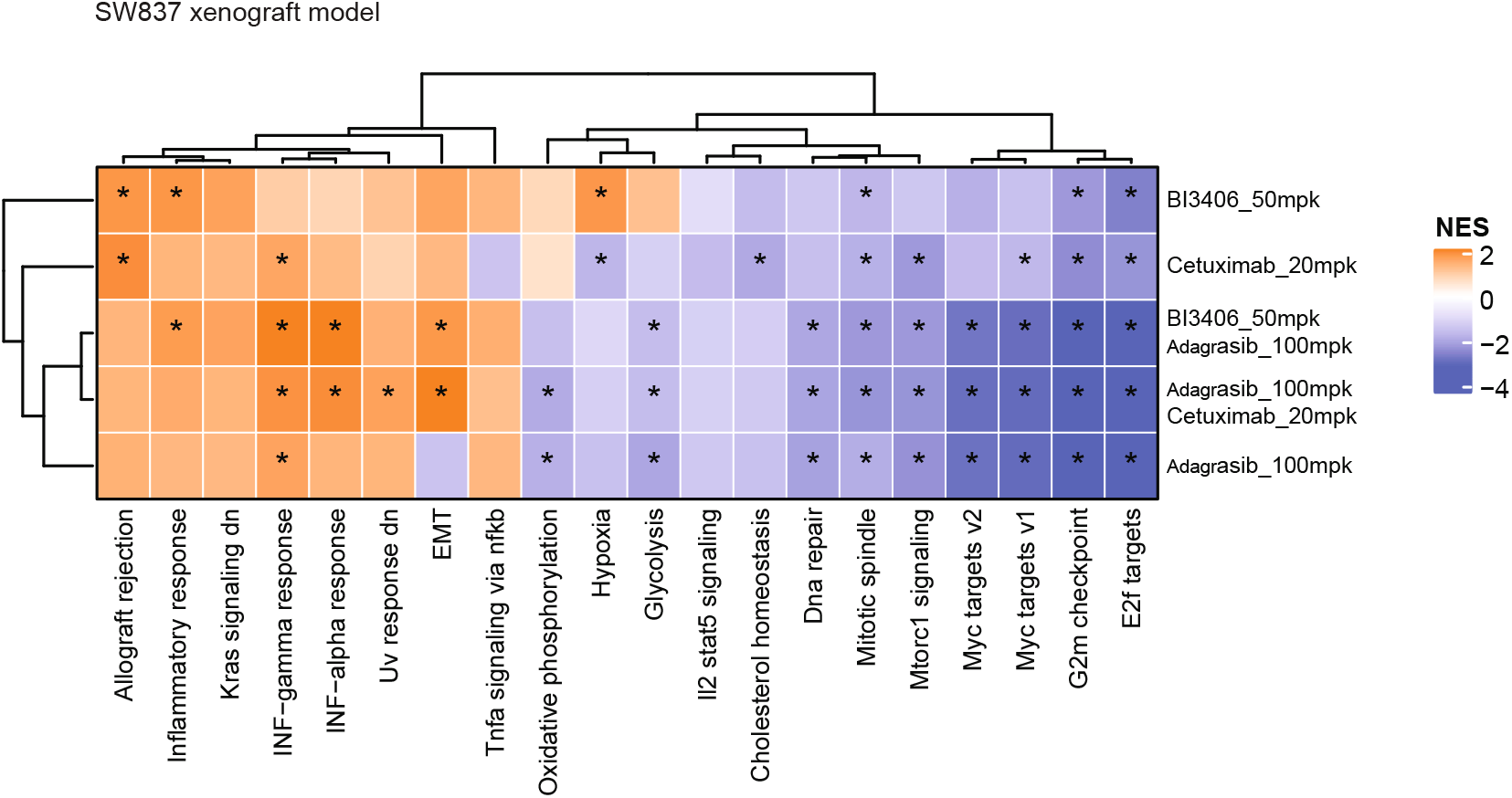
Differentially modulated pathways in a KRASG12C–driven colorectal cancer (CRC) xenograft model. Heatmap showing the normalized enrichment score of hallmark pathway gene sets in tumors from SW837 xenograft models treated with indicated single or combination therapies at 4 hours post-last dose.

**Extended Data Figure 3.**
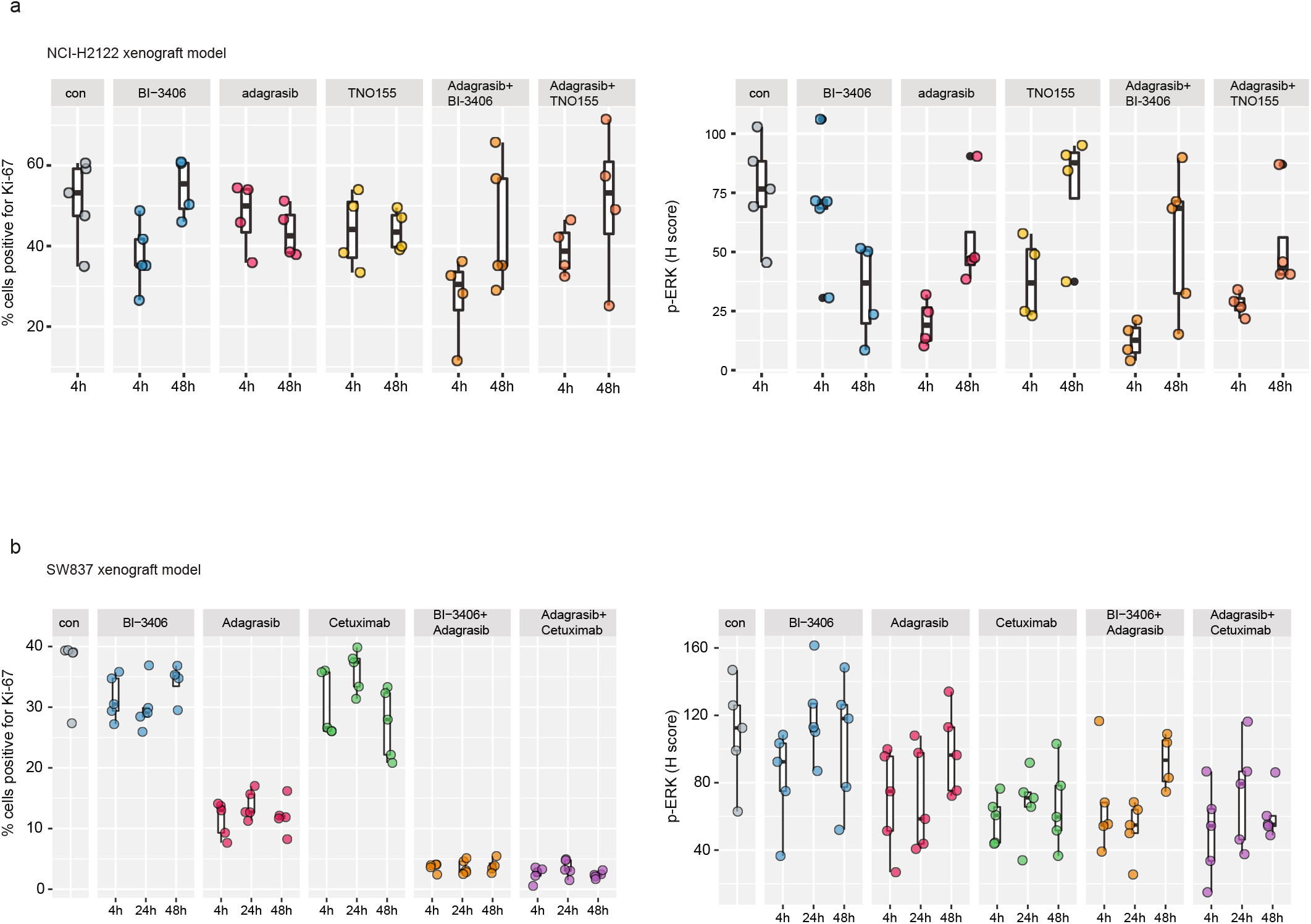
Differential gene expression induced by single-agent or combination treatments in a KRAS^G12C^–driven non-small cell lung cancer (NSCLC) or KRAS^G12C^–driven colorectal cancer (CRC) xenograft model. All treatments were administered over 7 consecutive days, with samples collected after the last dose. **a**, Biomarker analyses of cell proliferation biomarker, Ki-67 (left), and cell growth biomarker, phospho-ERK (p-ERK) (right), in NCI-H2122 xenograft models treated with vehicle (con), BI-3406 (50 mg/kg, bid, p.o.), adagrasib (100 mg/kg, qd, p.o.), TNO155 (10 mg/kg, bid, p.o.), adagrasib (100 mg/kg) plus BI-3406 (50 mg/kg), adagrasib (100 mg/kg) plus TNO155 (10 mg/kg) at 4 or 48 hours post-last dose. Control animals were treated with vehicle. N=4-5/group. Boxplots show low and upper quartiles and median line is indicates. Whiskers: 1.5 × interquartile range. Data analyzed by one-sided Mann-Whitney-Wilcoxon U-tests adjusted for multiple comparisons according to the Bonferroni-Holm Method within each subtopic. **b**, Biomarker analyses of cell proliferation biomarker, Ki-67 (left), and cell growth biomarker, phospho-ERK (p-ERK) (right), in SW837 xenograft models treated with vehicle (con), BI-3406 (50 mg/kg, bid, p.o.), adagrasib (100 mg/kg, qd, p.o.), TNO155 (10 mg/kg, bid, p.o.), adagrasib (100 mg/kg) plus BI-3406 (50 mg/kg), adagrasib (100 mg/kg) plus TNO155 (10 mg/kg) at 4 or 48 hours post-last dose. Control animals were treated with vehicle. N=4-5/group. Boxplots show low and upper quartiles and median line is indicates. Whiskers: 1.5 × interquartile range. Data analyzed by one-sided Mann-Whitney-Wilcoxon U-tests adjusted for multiple comparisons according to the Bonferroni-Holm Method within each subtopic.

**Extended Data Figure 4.**
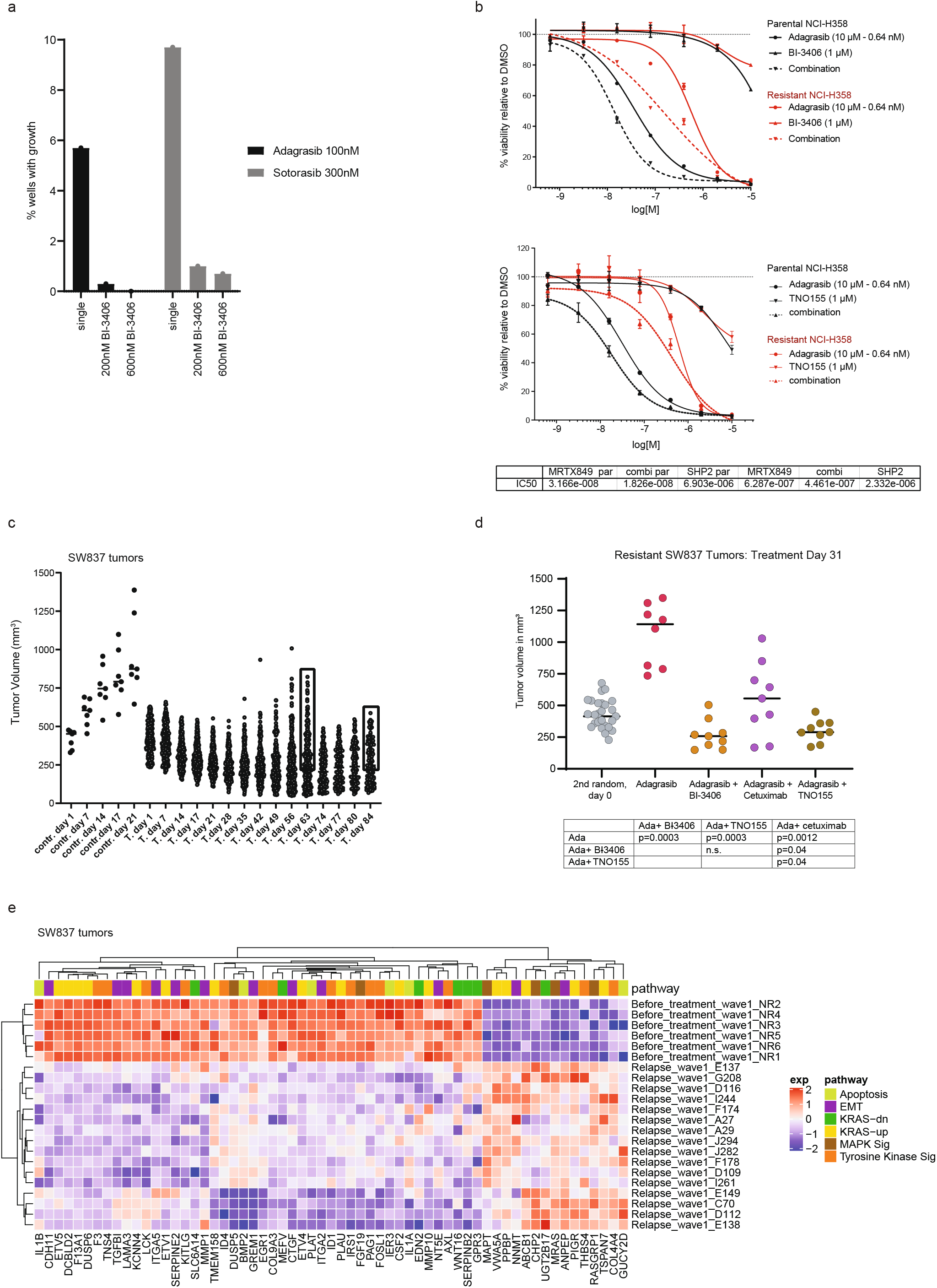
Combination treatments lead to regression in KRAS^G12C^i-resistant colorectal cancer (CRC) models. **a**, Growth response of BaF/3 cells harboring a KRAS^G12C^ single site variant library to KRAS^G12C^-specific inhibitors, adagrasib or sotorasib alone (single) or combined with 200 nM or 600 nM of BI-3406. N=1. **b**, Effects of adagrasib, BI-3406, or adagrasib plus BI-3406 (*top*) or adagrasib, TNO155, or adagrasib plus TNO155 (*bottom*) in a cell proliferation assay in adagrasib-sensitive (parental) or adagrasib-resistant NCI-H358 cells. Mean percentages of cell viability relative to the DMSO-treated control are shown. Error bars = ±SD; n=3 technical replicates. Each graph is representative of at least 2 independent experiments. **c**, Growth of SW837 tumors treated long-term with adagrasib (50mg/kg; n=300 mice total) or vehicle (0.5% Natrosol + 0.5% DMSO; n=7) on a 5 days on/2 days off dosing schedule. Outgrowth indicating tumors that developed acquired resistance to adagrasib is indicated with a black rectangle. **d**, Growth of adagrasib-resistant tumors per treatment group from (c) that were further treated with the following (n=5/group) for 31 days: adagrasib (50 mg/kg, qd, p.o.), adagrasib (50 mg/kg, qd, p.o.) plus BI-3406 (50mg/kg, bid, p.o.), adagrasib (50 mg/kg, qd., p.o.) plus cetuximab (20mg/kg, q3 or 4d, intraperitoneal), or adagrasib (50 mg/kg, qd) plus TNO155 (10mg/kg, bid). Bar indicates mean. Data were analyzed on Day 31 by an one-sided non-parametric Mann-Whitney-Wilcoxon U-tests adjusted for multiple comparisons according to the Bonferroni-Holm Method within each subtopic; ada = adagrasib; n.s. = non-significant. **e**, Heatmap showing expression z-score of genes belonging to apoptosis, epithelial-mesenchymal transition (EMT), or KRAS gene sets, as well as MAPK or tyrosine kinase signaling pathways from tumors collected from SW837 xenograft models prior to adagrasib treatment (Before Treatment) and during relapse (Relapse).

